## Appendix1 for "Misclassification in memory modification in *App^NL-G-F^* knock-in mouse model of Alzheimer’s disease"

### *Appendix 1*

#### *Alternative models*

In order to test whether the latent cause model (LCM) has better explanatory power in this study, we compared its fit to behavioral data with that of two alternative models: the variant of the Rescorla-Wagner (RW; Rescorla & Wagner, 1972) model and the latent state model (LSM; Cochran & Cisler, 2019). The RW model serves as a baseline, given its known limitations in explaining fear return after extinction, while the LSM is another contemporary model capable of modeling memory modification.

### Rescorla-Wagner (RW) model

The RW model, proposed by Rescorla and Wagner (1972), is an elemental theory in which an association between a compound of stimuli and an outcome consists of associations between individual stimulus and the outcome (Soto et al., 2014). In the present reinstatement experiment, the gross associative strength $V$ in each trial given the CS and the context is the sum of $V_{\mathrm{CS}}$ and $V_{\mathrm{context}}$:

$$V = V_{\mathrm{CS}} + V_{\mathrm{context}},$$

where the former is the associative strength between the CS and the US, while the latter is that between the context and the US. According to the original RW model, $V_{\mathrm{CS}}$ and $V_{\mathrm{context}}$ are updated as

$$\Delta V_{\mathrm{CS}} = \alpha_{\mathrm{CS}}\beta(\lambda - V),$$

$$\Delta V_{\mathrm{context}} = \alpha_{\mathrm{context}}\beta(\lambda - V),$$

where $\alpha$ and $\beta$ are the saliency of each stimulus and the learning rate for the US, respectively. $\lambda$ is the observed US that is the asymptote of $V$, and $(\lambda - V)$ expresses a prediction error between the observed US and $V$ as a predicted US given the stimuli. Consequently, $\Delta V_{CS}$ and $\Delta V_{context}$ would gradually increase during acquisition and decrease during extinction. However, in our reinstatement paradigm, this update rule causes an increase in $\Delta V_{CS}$ in the unsignaled shock trial, even though CS is not presented. In contrast, $\Delta V_{context}$ is underestimated due to the $V_{CS}$ in the prediction error term. To prevent this, we modified each update rule as

$$\Delta V_{\mathrm{CS}} = \alpha_{\mathrm{CS}}\beta(\lambda - V_{\mathrm{CS}})x_{\mathrm{CS}},$$

$$\Delta V_{\mathrm{context}} = \alpha_{\mathrm{context}}\beta(\lambda - V_{\mathrm{context}})x_{\mathrm{context}},$$

where $x_{\mathrm{CS}}$ and $x_{\mathrm{context}}$ were stimulus inputs while $\lambda$ was the US input, and these values were the same as those in the simulation in the latent cause model (see Material and methods). Thus, $V_{\mathrm{CS}}$ and $V_{\mathrm{context}}$ are updated only in the presence of the CS and the context, respectively. Finally, we estimated CR as a linear combination of stimulus $\mathbf{x}$ and associative strength $\mathbf{V}$,

$$\mathrm{CR} = x_{\mathrm{CS}}V_{\mathrm{CS}} + x_{\mathrm{context}}V_{\mathrm{context}}.$$

#### Latent state model (LSM)

The LSM, proposed by Cochran and Cisler (2019), shares several concepts and processes with the latent cause model (Gershman et al., 2017). In the LSM, an agent has an internal model such that $R(t)$, the outcome (e.g. US) at trial $t$, is generated according to an $n$-dimensional cue vector $\mathbf{c}(t)$ (e.g. CS), $l$-th latent state in a total of $L$ latent states acquired so far, and a latent noise. The series of latent states across trials is a Markov chain, as each latent state changes at probability $\gamma(L - 1) / L$, otherwise stays, where $\gamma$ is a parameter that governs the transition probability of latent states. $R(t)$ given $l$-th latent state is approximated by a linear combination of $\mathbf{c}$ and an associative strength vector $\mathbf{v}$ as the RW model, resulting in the prediction error $E_l(t) = R(t) - \mathbf{c}(t)'\mathbf{v}_l(t)$. $\mathbf{v}$ is iteratively updated by the "rumination" process at the end of this trial (see below). As in the latent cause model, the purpose of its computation is to minimize the prediction error.

At the beginning of each trial, the agent observes $\mathbf{c}(t)$ and $R(t)$, then calculates the posterior probability of each latent state $p_l(t)$ by Bayes' rule,

$$p_l(t) = \frac{1}{l_0}\left((1-\gamma)p_l(t-1) + \frac{\gamma}{L}\right)\phi\left(\frac{E_l(t)}{\sigma(t)}\right)$$

where $\varphi(\cdot)$ is the likelihood of the prediction error given latent state $l$, evaluated using a standard normal distribution. $\sigma(t)$ represents the uncertainty of $R(t)$, and is updated over trials based on the prediction errors (see below). The term before the likelihood is the prior of latent state $l$, and $l_0$ is the partition function, the sum over $p_l(t)$ across latent states 1 to $L$. Next, the agent determines whether a new latent state should be inferred using the change point statistic $q(t)$,

$$q(t+1) = max\left(q(t) + \log\left(\frac{\phi(0)}{l_0}\right) - \delta, 0\right)$$

where $\varphi(0)$ represents the ideal state with no prediction error, hence $\log(\varphi(0) / l_0)$ can be interpreted as the log likelihood ratio between this ideal state and the gross likelihood provided by $L$ latent states. $\delta$ is the penalty term. Intuitively, $q(t)$ indicates how the set of latent states acquired so far is close to the ideal state. If $q(t)$ exceeds the threshold $\eta$, $L$+1-th new state is inferred while $L < L_{max}$, where $L_{max}$ is the predefined maximum number of latent states that can be acquired.

Given $\mathbf{c}(t)$ and $E_l(t)$, associative strength $\mathbf{v}$ of latent state $l$ is updated a

$$\mathbf{v}_l(t+1) = \mathbf{v}_l(t) + \mathbf{A}_l(t)\boldsymbol{c}(t)E_l(t)$$

where $\mathbf{A}_l(t)$ is the associability matrix for latent state $l$. $\mathbf{A}_l(t)$ can be viewed as a learning rate as it governs how quickly associative strength changes. It depends on the probability of latent state $l$ and the history of cue observations, according to the effort matrix $\mathbf{B}_l(t)$ under latent state $l$, that is,

$$\mathbf{A}_l(t) = \alpha_0 p_l(t)\mathbf{B}_l(t)^{-1}$$

and $\mathbf{B}_l(t)$ is updated by

$$\mathbf{B}_l(t+1) = \mathbf{B}_l(t) + \alpha_2(p_l(t)\boldsymbol{c}(t)\boldsymbol{c}(t)' - \mathbf{B}_l(t))$$

where $\alpha_0$ and $\alpha_2$ are the learning rate and ranging between 0 and 1. Then the agent updates uncertainty $\sigma(t)$ of $R(t)$ by

$$\sigma^2(t+1) = \sigma^2(t) + \alpha_1(\mathbb{E}(E^2(t)) - \sigma^2(t))$$

where $\alpha_1$ is the learning rate for the uncertainty ranging between 0 and 1. $\mathbb{E}(E^2(t))$ is the squared prediction error averaged over latent states. The initial value of $\sigma^2$, denoted as $\sigma_0^2$, is predefined. By the beginning of next trial $t + 1$, $p_l(t)$ is updated by

$$p_l(t)^* = (1-\gamma)^{ITI-1} p_l(t) + (1-(1-\gamma)^{ITI-1})\frac{1}{L}$$

expressing the temporal decay over inter-trial interval ($ITI$) and then used as the prior in the next trial. In addition, the "rumination" process runs given the ITI: the agent iteratively updates $\mathbf{v}_l$ using the same observation of $\mathbf{c}(t)$ and $R(t)$ from the previous trial. The number of this iteration is determined by min ($\chi$, $ITI$-1), where $\chi$ is predefined.

### Evaluation of model fit

#### Settings

To compare the model fit among the RW model, the LSM, and the LCM, we initially estimated the parameters of models minimizing the squared prediction errors between the observed CR and the simulated CR given the proposed parameters. To apply the same evaluation criteria for prediction error set in the LCM, we normalized both the observed CR and the simulated CR in the RW model and the LSM.

For the RW model, we estimated $\alpha_{CS}$, $\alpha_{context}$, and $\beta$ using the least squares error method in the grid search. We assumed that the value of each parameter could take any one of a uniform grid of 50 points between 0.1 and 1, therefore there were $50^3$ possible combinations of the 3 parameters. For each observed CR, we selected the combination of parameters with the lowest sum of squared prediction errors across all trials. The criteria for anomaly detection were the same as those for the LCM, except that the Gelman-Rubin statistic was not applied (see Material and Method).

For the LSM, we estimated $\alpha_0$, $\alpha_1$, $\alpha_2$, $\gamma$, $L_{\max}$, $\eta$, $\chi$ and $\sigma_0$, but fixed two parameters, $\tau$ (exploration-exploitation parameter) and $\delta$ (constant involved in change point statistic), assuming that their effects would be minimal. $\tau$ and $\delta$ were set to the default value in Cochran and Cisler (2019). We simulated the reinstatement paradigm given the proposed parameters, using the codes provided by Cochran and Cisler (2019) (https://github.com/cochran4/OnlineLatentStateLearning.git). Note that the variable names above follow those in the code, except $L_{\max}$; $L_{\max}$ is noted *ncop* in the code; $\alpha_1$, $\alpha_2$ and $\eta$ in the code correspond to $\alpha_0$, $\beta_0$ and $v$ in Table 3 of Cochran and Cisler (2019). The values of the CS, context, US, *ITI*, and other experimental designs were the same as those set in parameters estimation in the LCM (see Material and Method), whereas the initial bias for the US centering was set to 0. We treated the expected outcome given cues, $\mathbf{c}(t)'\mathbf{v}(t)$, as the CR at each trial. As in the LCM, the proposed parameters were sampled while they reduced the prediction errors, through the MCMC method with the slice sampling (see Material and Method). The step size in *slicesample* function was changed from 100 to 50 to reduce the sampling time. As only three samples in the 12-month-old group were marked as anomalies due to the failure of the convergence, not prediction error, the quality of sampled parameters should be comparable to those of the LCM, in terms of convergence. The initial guess, upper bound, and lower bound of each parameter are shown in Appendix 1 – table 1. The criteria for anomaly detection were the same as those for the LCM (see Material and Method).

#### Simulation

To illustrate the representative fitting characteristics of each model, we initially estimated parameters from the median CR curves in each group. The RW model given the estimated parameters successfully replicated the gradual increase of CRs in the acquisition and their decrease in the extinction (Appendix 1 – figure 1A to 1D). As expected, the RW model failed to replicate the reinstatement, as a single trial of context-US pairing, the unsignaled shock, only resulted in a small increase in the overall associative strength after sufficient extinction. Indeed, only the simulated CR of 12-month-old $App^{NL\text{-}G\text{-}F}$ mice groups passed the anomaly detection test. In the LSM, the simulated CR well replicated the gradual increase of CR in the acquisition in the 6-month-old group than in the 12-month-group, while the gradual decrease of CR was better reproduced in the 12-month-group than in the 6-month-old group (Appendix 1 – figure 1E to 1H). The increased CR in the first trial in the second extinction phase (trial 24) was captured only in the 12-month-old control (Appendix 1 – figure 1G). The magnitude of reinstatement was well replicated in all groups other than the 6-month-old control (Appendix 1 – figure 1E-H).

The number of samples classified as anomalies in each group was highest in the RW model across all groups, and that in the LSM was twice that of in the LCM except 6-month-old $App^{NL\text{-}G\text{-}F}$ mice, suggesting that the LCM was robust for individual differences (Appendix 1 – figure 2). The prediction errors of all accepted samples in the RW model were obviously higher in the reinstatement at trial 36 (Appendix 1 – figure 3A), which is aligned with the median data (Appendix 1 – figure 1A-D). In contrast, the prediction errors of accepted samples in the LSM at trial 36 were smaller (Appendix 1 –

figure 3B), suggesting that it could replicate the reinstatement, as shown in the median data (Appendix 1 – figure 1E to 1H). In both models, the prediction errors during the extinction phase were lower than in other phases (Appendix 1 – figure 3A and 3B). We then statistically compared the sum of prediction errors over trials, as well as those in acquisition trials and test phase trials, between the LCM and LSM, excluding the RW model due to the lower number of accepted samples (Appendix 1 – figure 2). The sum of prediction error over trials was significantly higher in the LCM, especially in the 12-month-old group (Appendix 1 – figure 4A and 4B, left panel). This would be due to the significantly larger prediction errors during the acquisition phase in the LCM (Appendix 1 – figure 4A and 4B, center panel). Notably, the prediction errors in the 3 test phases were almost comparable (Appendix 1 – figure 4A and 4B, right panel).

In this study, we favored models that robustly replicate CR during the 3 test phases, rather than during the acquisition phase, from as many samples as possible. In this sense, the LSM remains a potential candidate as it successfully replicated the reinstatement in a different manner from that in the LCM. There were only one or two latent states inferred during the acquisition (Appendix 1 – figure 5A and 5D, left panel). Almost all accepted samples continued to infer the acquisition latent states during extinction, without inferring new states (Appendix 1 – figure 5B and 5E, left panel). Although a new state was inferred after extinction in some of the samples, its posterior probability was low in test 3 (Appendix 1 – figure 5C and 5F). On the contrary, the sum of posterior probabilities of acquisition latent states was highest in test 3 (Appendix 1 – figure 5A and 5D), suggesting that the reinstatement is explained by the return of the acquisition state. However, this pattern was also observed in the 12-month-old $App^{NL\text{-}G\text{-}F}$ group. When we calculated the DIs between tests from simulated CR in the LSM (Appendix 1 – figure 6), the significantly lower DI between test 3 and test 1 in the 12-month-old $App^{NL\text{-}G\text{-}F}$ group (Figure 2H) was indeed not replicated (Appendix 1 – figure 6C), indicating accepted samples in the LSM failed to capture our behavioral features of interest.

Thus, we conclude that standing on the LCM to explain the memory modification observed in this study would be reasonable, given its robustness for individual differences and the reproducibility of the empirical data of interest. On the other hand, we did not test all possible combinations of parameters of the LSM or all possible experimental protocols to induce reinstatement. Since the conditions for simulating reinstatement and estimating parameters were initially developed in the LCM and applied *post hoc* to the LSM, we did not rule out the possibility that the LSM can demonstrate better explanatory power than the LCM in alternative conditions. A comprehensive search of such conditions may be a valuable direction for future research.

.

**Appendix 1 – table 1. The initial value, lower bound, and upper bound of parameters in slice sampling in the latent state model**

| Description | Parameters | Initial value | Lower bound | Upper bound |
|---|---|---|---|---|
| learning rate for associative strength | $\alpha_0$ | 0.05 | 0.005 | 0.1 |
| learning rate for variance | $\alpha_1$ | 0.06 | 0.005 | 0.5 |
| learning rate for covariance | $\alpha_2$ | 0.04 | 0.005 | 0.5 |
| transition probability between states | $\gamma$ | 0.1 | 0.01 | 1 |
| maximum number of latent states | *ncop* | 15 | 1 | 36 |
| threshold to activate new state | $\eta$ | 2.5 | 0.01 | 3.05 |
| number of rumination updates | $\chi$ | 5 | 1 | 10 |
| initial standard deviation | $\sigma_0$ | 0.5 | 0.1 | 1 |

*Note.* $\tau$ and $\delta$ were fixed at 10 and 0.6, respectively, in the present simulation.

**Appendix 1 – figure 1. Model fit of the Rescorla-Wagner model and the latent state model to the behavioral data.**

**(A)** Simulation of reinstatement in the 6-month-old control mice in the RW model. The estimated parameter value: $\alpha_{CS} = 0.1$, $\alpha_{context} = 1$, $\beta = 0.504$. **(B)** Simulation of reinstatement in the 6-month-old *App*$^{NL\text{-}G\text{-}F}$ mice in the RW model. The estimated parameter value: $\alpha_{CS} = 0.1$, $\alpha_{context} = 1$, $\beta = 0.302$. **(C)** Simulation of reinstatement in the 12-month-old control mice in the RW model. The estimated parameter value: $\alpha_{CS} = 0.1$, $\alpha_{context} = 1$, $\beta = 0.614$. **(D)** Simulation of reinstatement in the 12-month-old *App*$^{NL\text{-}G\text{-}F}$ mice in the RW model. The estimated parameter value: $\alpha_{CS} = 0.1$, $\alpha_{context} = 1$, $\beta = 0.596$. **(E)** Simulation of reinstatement in the 6-month-old control mice in the LSM. The estimated parameter value: $\alpha_0 = 0.012$, $\alpha_1 = 0.020$, $\alpha_2 = 0.040$, $\gamma = 0.020$, ncop = 10, $\eta = 2.369$, $\chi = 3$, $\sigma_0 = 0.99$. **(F)** Simulation of reinstatement in the 6-month-old *App*$^{NL\text{-}G\text{-}F}$ mice in the LSM. The estimated parameter value: $\alpha_0 = 0.078$, $\alpha_1 = 0.058$, $\alpha_2 = 0.418$, $\gamma = 0.258$, ncop = 10, $\eta = 2.290$, $\chi = 7$, $\sigma_0 = 0.69$. **(G)** Simulation of reinstatement in the 12-month-old control mice in the LSM. The estimated parameter value: $\alpha_0 = 0.043$, $\alpha_1 = 0.191$, $\alpha_2 = 0.376$, $\gamma = 0.099$, ncop = 10, $\eta = 0.044$, $\chi = 8$, $\sigma_0 = 0.58$. **(H)** Simulation of reinstatement in the 12-month-old *App*$^{NL\text{-}G\text{-}F}$ mice in the LSM. The estimated parameter value: $\alpha_0 = 0.069$, $\alpha_1 = 0.020$, $\alpha_2 = 0.495$, $\gamma = 0.886$, ncop = 32, $\eta = 0.040$, $\chi = 9$, $\sigma_0 = 0.62$.

**(A to H)** The observed CR is the median freezing rate during the CS presentation over the mice within each group; both observed and simulated CR of each group were divided by their maximum over all trials. The vertical dashed lines indicate the boundaries of phases. In the RW model, 6-month-old control (A), 6-month-old *App*$^{NL\text{-}G\text{-}F}$ (B), and 12-month-old control (C) were marked as anomalies in parameter estimation.

**Appendix 1 – figure 2. Number of excluded samples during parameter estimation in each model.** The number of samples that did not pass the accepted criteria in the 6-month-old group **(A)** and 12-month-old group **(B)**. LCM: latent cause model; LSM: latent state model; RW: Rescorla-Wagner model. Colors indicate different groups: orange represents 6-month-old control ($n$ = 24), light blue represents 6-month-old $App^{NL\text{-}G\text{-}F}$ mice ($n$ = 25), pink represents 12-month-old control ($n$ = 24), and dark blue represents 12-month-old $App^{NL\text{-}G\text{-}F}$ mice ($n$ = 25). The transparency of the color indicates accepted samples (dark) and anomalies (light).

**Appendix 1 – figure 3. The squared prediction error in the Rescorla-Wagner model (A) and the latent state model (B).**
The squared prediction error across trials is shown as median with interquartile range. In the RW model, data from all samples are used. In the LSM, only accepted samples are shown (the exact numbers are shown in Appendix 1 – figure 2).

**Appendix 1 – figure 4. Comparison of squared prediction error among models in the 6-month-old group (A) and 12-month-old group (B).**
The data are shown as median with interquartile range. In the RW model, data from all samples are used. In the LSM and LCM, only accepted samples are shown (the exact numbers are shown in Appendix 1 – figure 2). LCM: latent cause model; LSM: latent state model; RW: Rescorla-Wagner model. *$p < 0.05$, **$p < 0.01$, and ***$p < 0.001$ by Mann-Whitney $U$ test comparing LCM and LSM. The RW model was excluded from the statistical comparison because of the low sample number.

**Appendix 1 – figure 5. The number of latent states inferred in different phases and their posterior probability in test 3.**
**(A to E)** The figure format is the same as that in Figures 4C-E and 5C-E for the latent cause model. The latent states were classified based on the trial they were initially inferred: during the acquisition trials (first row), extinction trials (second row), and trials after extinction (third row). The first and third columns show the count of latent states in the 6-month-old group and 12-month-old group, respectively. The second and fourth columns show the posterior probability of latent states in test 3 in the 6-month-old group and 12-month-old group, respectively. The horizontal axis indicates the proportion of the value in each group. Colors in the histogram indicate different groups: orange represents 6-month-old control ($n$ = 6), light blue represents 6-month-old $App^{NL\text{-}G\text{-}F}$ mice ($n$ = 13), pink represents 12-month-old control ($n$ = 10), and dark blue represents 12-month-old $App^{NL\text{-}G\text{-}F}$ mice ($n$ = 11). Each black dot represents one animal.

**Appendix 1 – figure 6. Discrimination index (DI) between test 2 and test 1 (A), between test 3 and test 2 (B), and between test 3 and test 1 (C) calculated from simulated CR in the latent state model.**

The dashed horizontal line indicates DI = 0.5, which means no discrimination between the two phases. $^{\dagger}p < 0.05$ by one-sample Student's *t*-test, and the alternative hypothesis specifies that the mean differs from 0.5; $*p < 0.05$ by one-way ANCOVA with age as a covariate; No significant difference between control and $App^{NL\text{-}G\text{-}F}$ mice within the same age was detected by Student's *t*-test comparing. Colors in the box plot indicate different groups: orange represents 6-month-old control ($n$ = 6), light blue represents 6-month-old $App^{NL\text{-}G\text{-}F}$ mice ($n$ = 13), pink represents 12-month-old control ($n$ = 10), and dark blue represents 12-month-old $App^{NL\text{-}G\text{-}F}$ mice ($n$ = 11). Each black dot represents one animal.

### Appendix 1 – figure 1

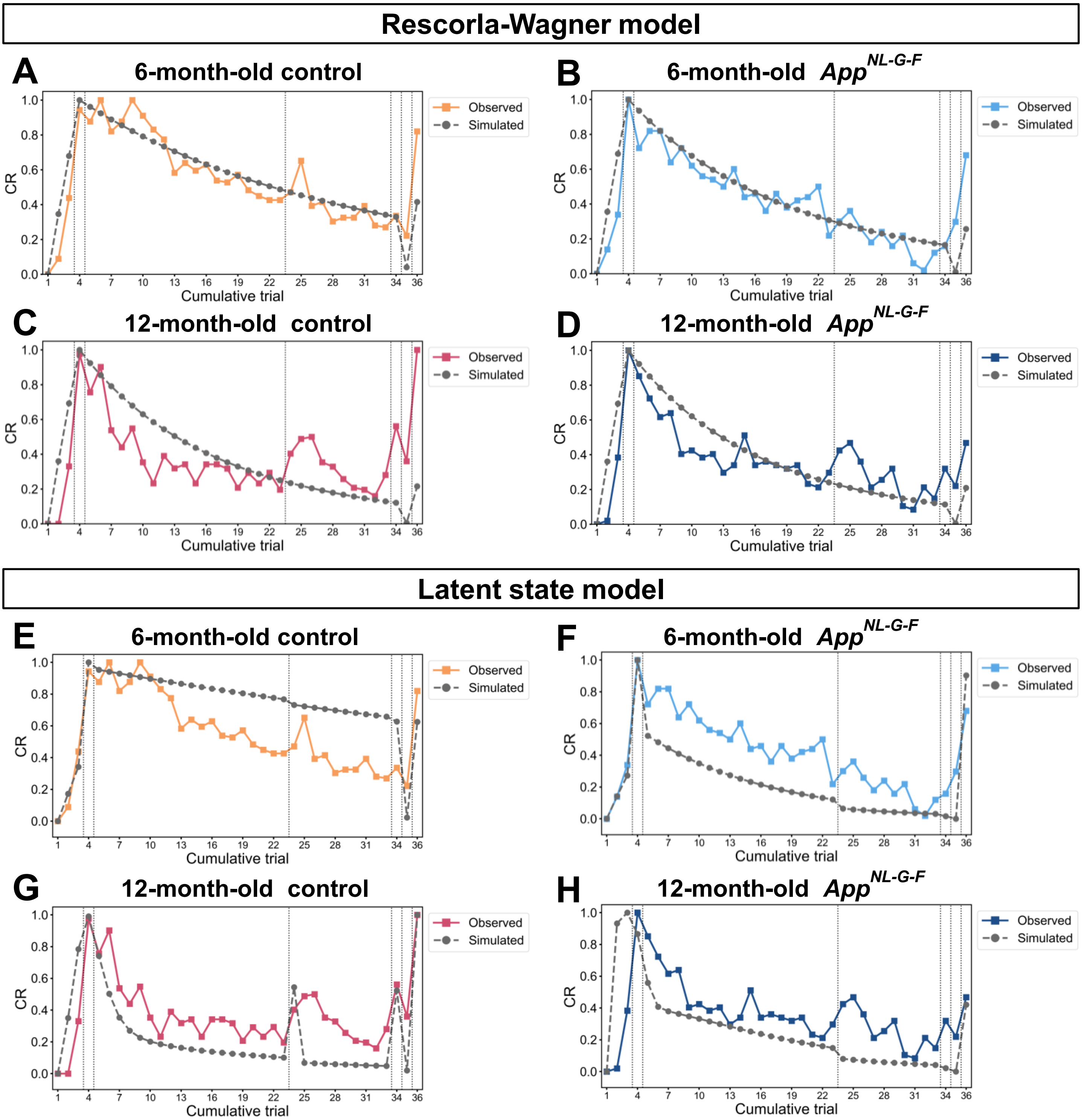

### Appendix 1 – figure 2

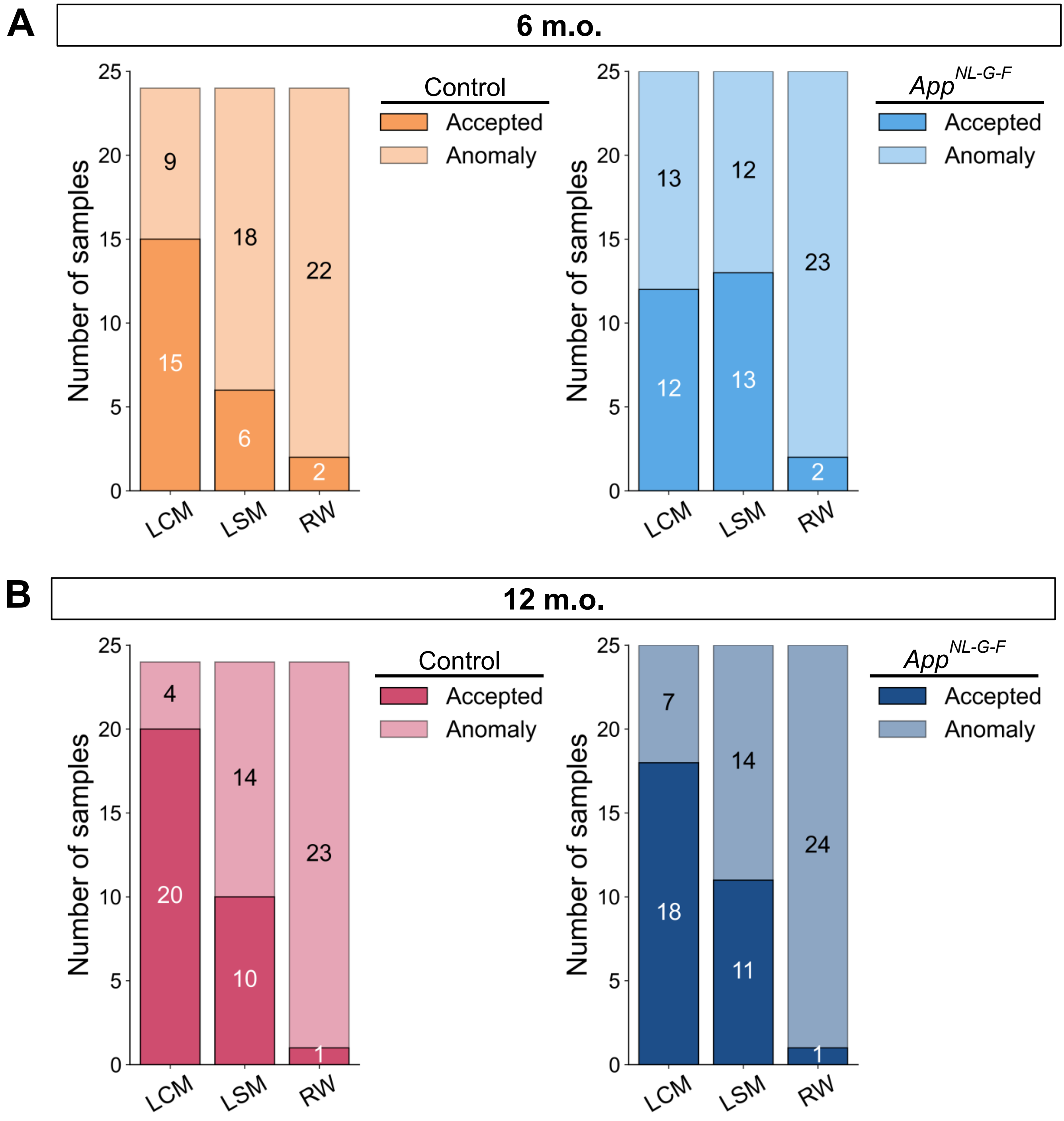

### Appendix 1 – figure 3

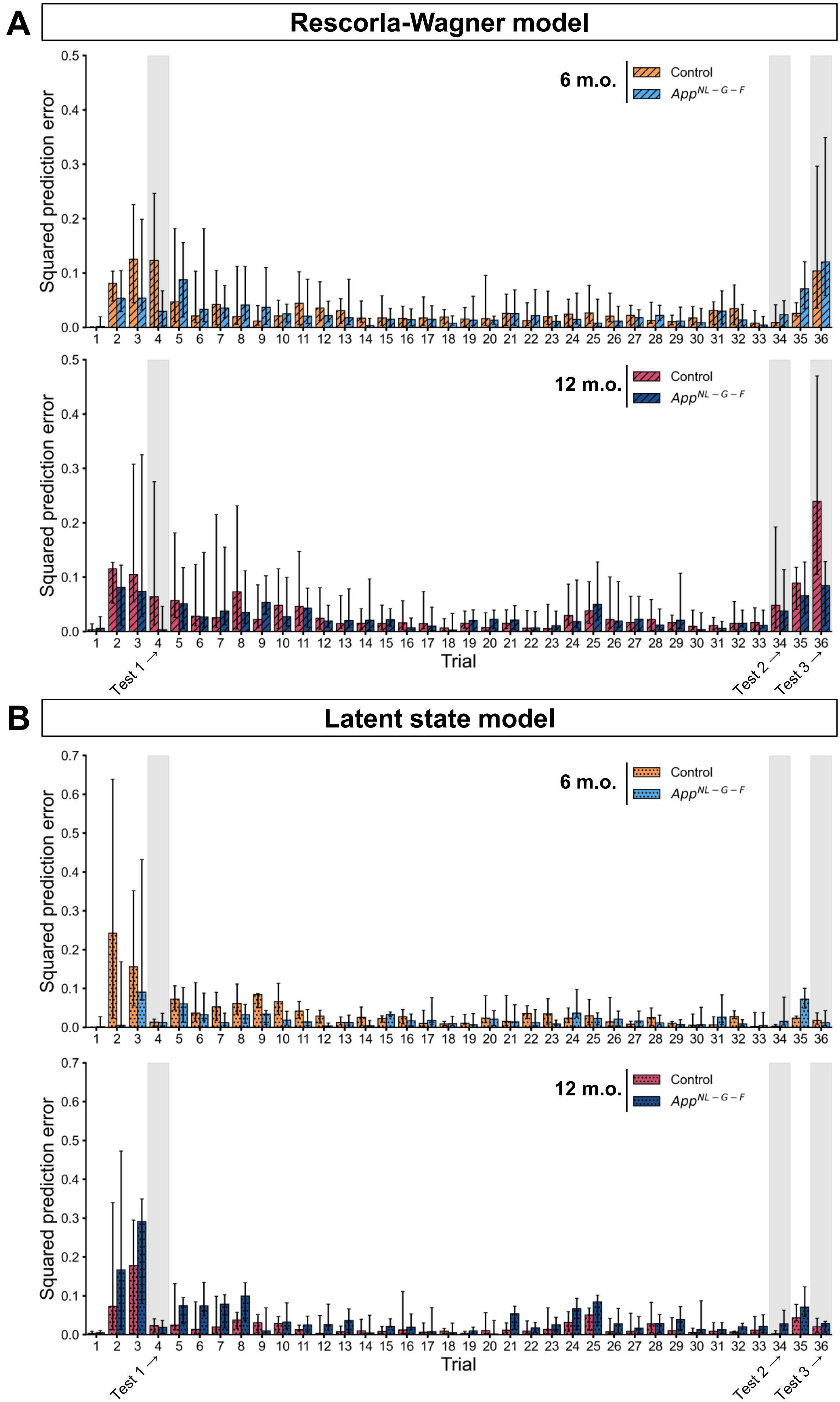

### Appendix 1 – figure 4

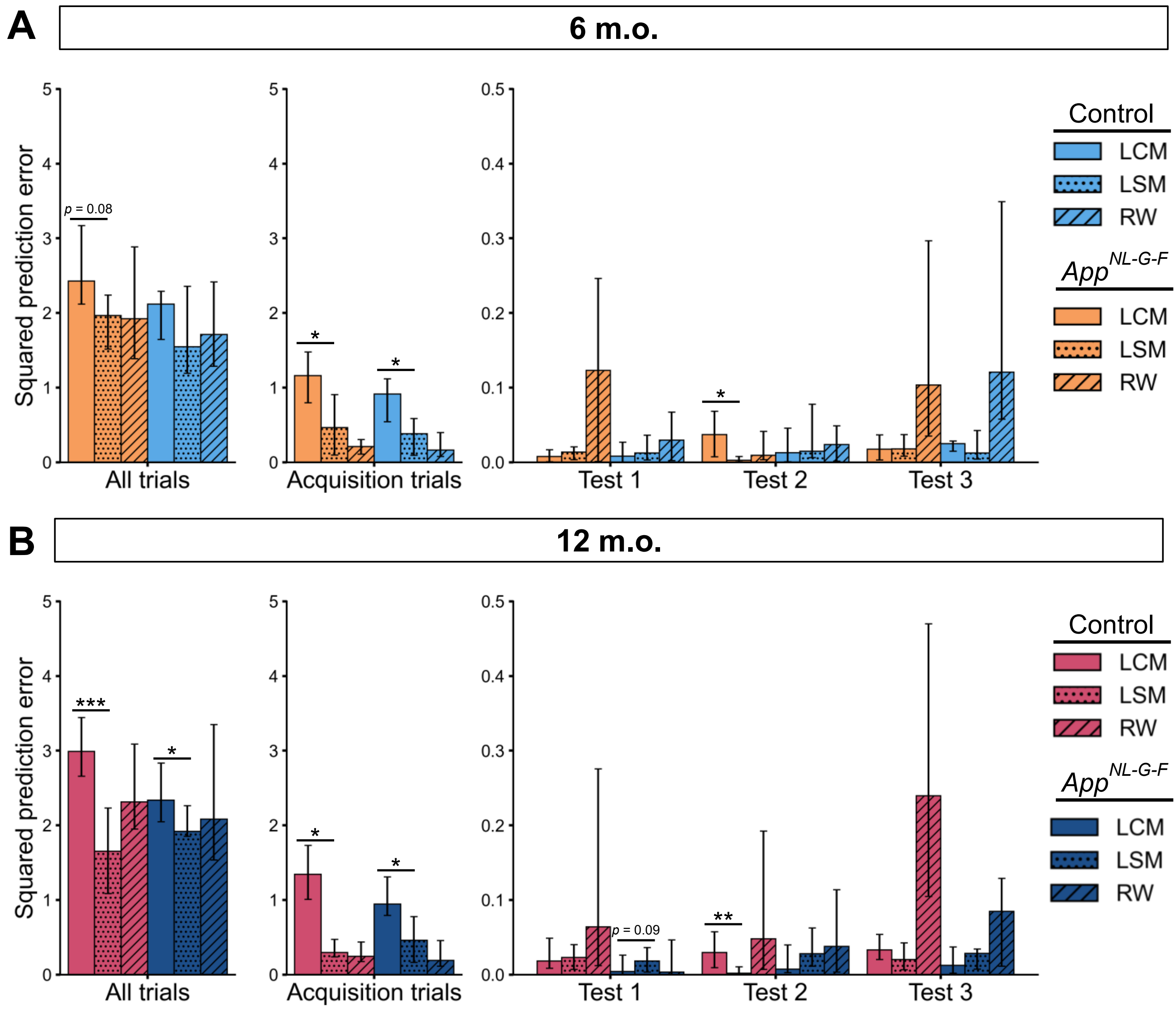

### Appendix 1 – figure 5

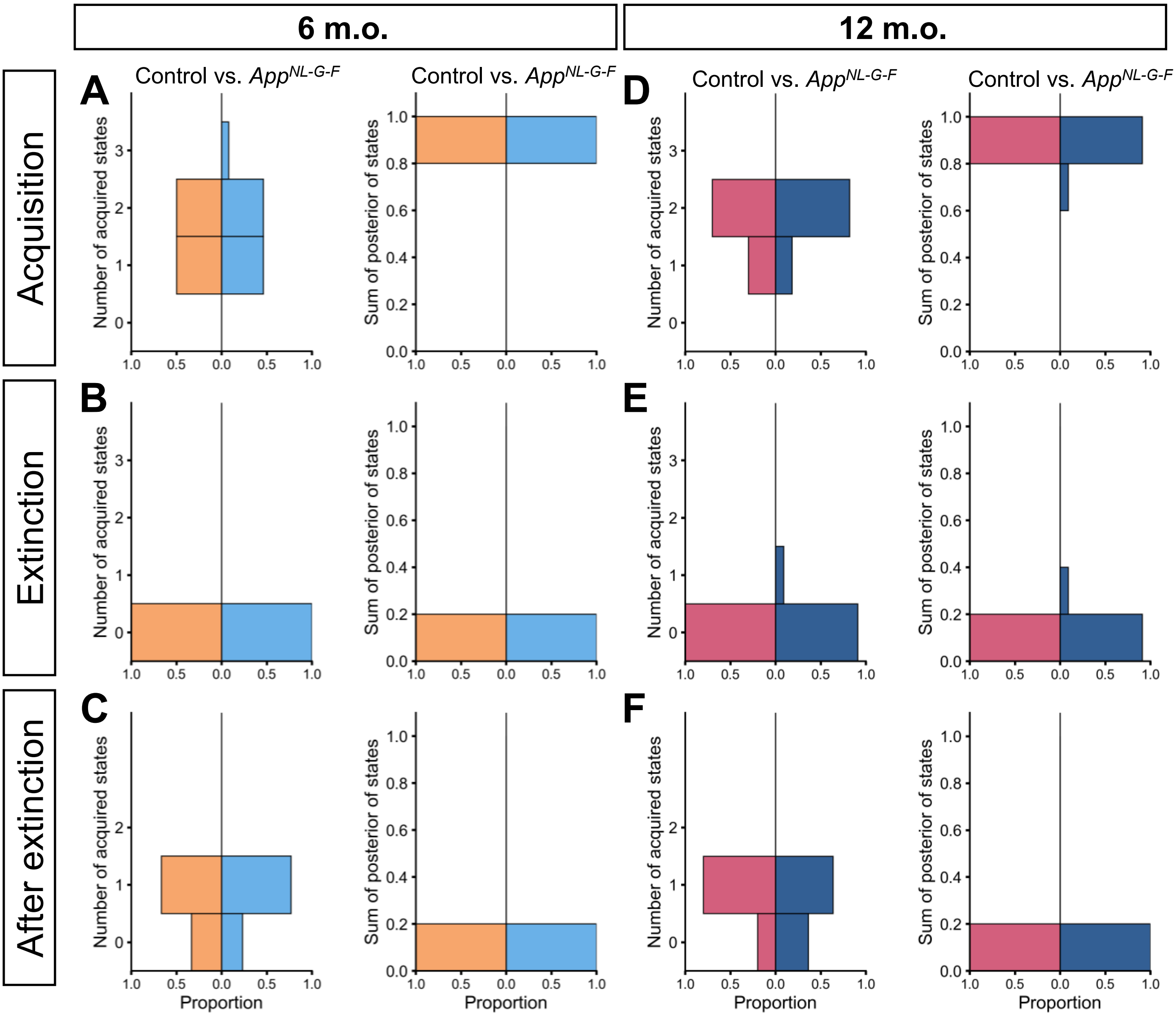

### Appendix 1 – figure 6

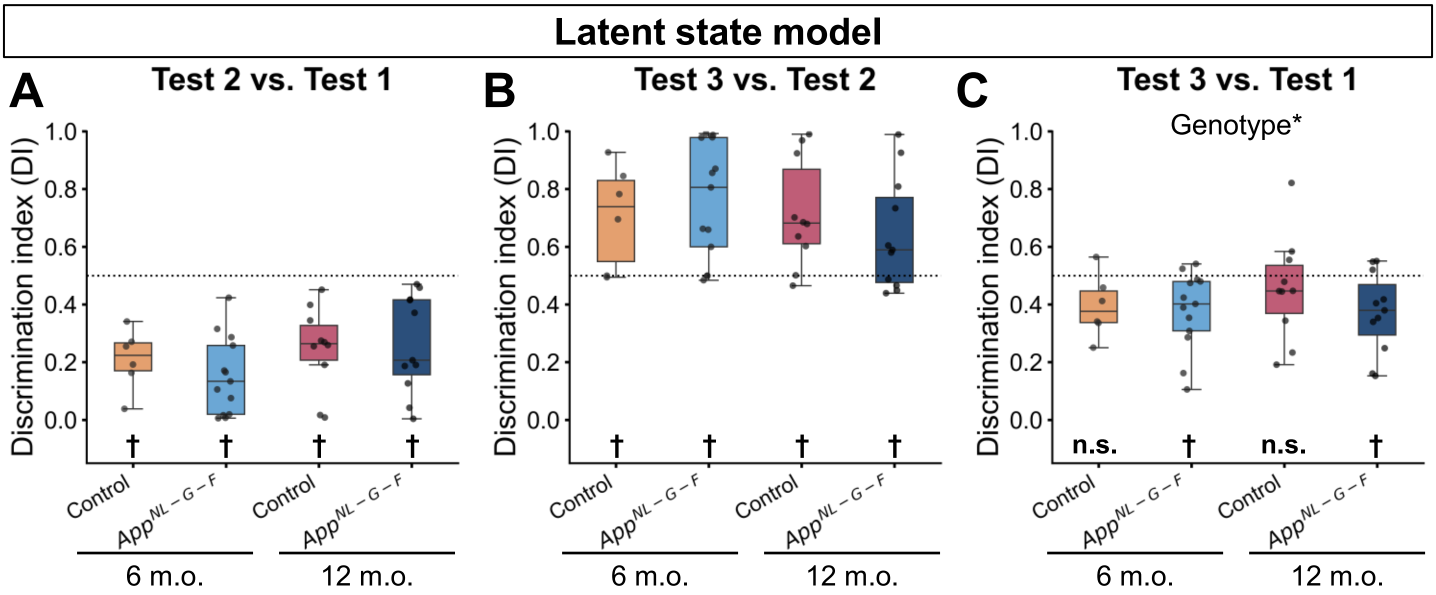
