## Appendix2 for "Misclassification in memory modification in *App^NL-G-F^* knock-in mouse model of Alzheimer’s disease"

### *Appendix 2*

#### *Relationship between parameters, conditioned response, and internal states*

**Appendix 2 – table 1. The initial value, lower bound, and upper bound of parameters in slice sampling**

| Parameters | Initial value | Lower bound | Upper bound |
|---|---|---|---|
| $\alpha$ | 1.5 | 1 | 3 |
| $g$ | 1 | 0.01 | 2 |
| $\eta$ | 0.1 | 0.01 | 1 |
| maxIter[a] | 3 | 1 | 5 |
| $w_0$ | 0 | -0.01 | 0.01 |
| $\sigma_r^2$ | 0.4 | 0.01 | 3 |
| $\sigma_x^2$ | 1 | 0.01 | 3 |
| $\theta$ | 0.01 | 0.002 | 0.02 |
| $\lambda$ | 0.01 | 0.002 | 0.02 |
| $K$ | 33 | 1 | 36 |

[a] max. no. of iterations.

**Appendix 2 – table 2. Effect of $\alpha$ and $\eta$ on test trials CR, number of latent causes, and test 3 internal states**

| $\alpha$ | $\eta$ | Test 1 | Test 2 | Test 3 | $K_{acq}$ | $K_{ext}$ | $K_{rem}$ | Test 3 | | | | | |
|---|---|---|---|---|---|---|---|---|---|---|---|---|---|
| | | | | | | | | $P_{acq}$ | $P_{ext}$ | $P_{rem}$ | $w_{context, acq}$ | $w_{context, ext}$ | $w_{context, rem}$ |
| 1 | 0.01 | 0.78 | 0.18 | 0.19 | 2 | 2 | 1 | 0.026 | 0.477 | 0.497 | 0.010 | 0.006 | 0.000 |
| 1 | 0.5 | 1.00 | 0.64 | 0.93 | 2 | 2 | 1 | 0.017 | 0.482 | 0.502 | 0.147 | 0.288 | 0.000 |
| 3 | 0.01 | 0.19 | 0.17 | 0.18 | 3 | 30 | 0 | 0.043 | 0.957 | 0.000 | 0.010 | 0.006 | 0.000 |
| 3 | 0.5 | 1.00 | 0.49 | 1.00 | 2 | 30 | 1 | 0.049 | 0.863 | 0.089 | 0.155 | 0.049 | 0.243 |

*Note*. This table is used for the reply to reviewer comment #14. The fixed parameter value: $g = 1$, max. no. of iteration = 3, $w_0 = 0$, $\sigma_r^2 = 0.4$, $\sigma_x^2 = 1$, $\theta = 0.01$, $\lambda = 0.01$, $K = 33$. Test 1, Test 2, Test 3: the simulated CR in each test. $K_{acq}$, $K_{ext}$, $K_{rem}$: the number of latent causes inferred during acquisition phases, extinction phases, and the remaining trials. $P_{acq}$, $P_{ext}$, $P_{rem}$: the posterior probability of latent causes inferred during acquisition phases, extinction phases, and the remaining trials in test 3. $w_{context, acq}$, $w_{context, ext}$, $w_{context, rem}$: the associative weight between context and US in the latent causes inferred during acquisition phases, extinction phases, and the remaining trials in test 3.

**Appendix 2 – table 3. Effect of $w_0$, $\theta$, and $\lambda$ on CR at each trial in the acquisition phase**

| $w_0$ | $\theta$ | $\lambda = 0.01$ | | | $\lambda = 0.02$ | | |
|---|---|---|---|---|---|---|---|
| | | Trial 1 | Trial 2 | Trial 3 | Trial 1 | Trial 2 | Trial 3 |
| -0.01 | 0.002 | 0.080757 | 0.999969 | 1 | 0.241964 | 0.977479 | 1 |
| 0 | 0.002 | 0.42074 | 1 | 1 | 0.460172 | 0.994947 | 1 |
| 0.01 | 0.002 | 0.841345 | 1 | 1 | 0.691462 | 0.999156 | 1 |
| -0.01 | 0.01 | 0.013903 | 0.999333 | 1 | 0.135666 | 0.945672 | 0.999999 |
| 0 | 0.01 | 0.158655 | 0.999993 | 1 | 0.308538 | 0.98508 | 1 |
| 0.01 | 0.01 | 0.57926 | 1 | 1 | 0.539828 | 0.996929 | 1 |
| -0.01 | 0.02 | 0.000687 | 0.986396 | 1 | 0.054799 | 0.865261 | 0.999994 |
| 0 | 0.02 | 0.02275 | 0.999588 | 1 | 0.158655 | 0.952757 | 1 |
| 0.01 | 0.02 | 0.211855 | 0.999996 | 1 | 0.344578 | 0.987459 | 1 |

*Note*. This table is used for the reply to reviewer comment #26. The fixed parameter value: $\alpha = 1.5$, $g = 1$, $\eta = 0.1$, max. no. of iteration = 3, $\sigma_r^2 = 0.4$, $\sigma_x^2 = 1$, $K = 33$

**Appendix 2 – table 4. Effect of $\alpha$ and $g$ on the number of latent causes inferred at each phase**

| $\alpha$ | $g$ | $K_{acq}$ | $K_{ext}$ | $K_{rem}$ |
|---|---|---|---|---|
| 1 | 0.01 | 2 | 0 | 0 |
| 2 | 0.01 | 2 | 0 | 0 |
| 1 | 1 | 2 | 2 | 1 |
| 2 | 1 | 2 | 30 | 1 |

*Note*. This table is used for the reply to reviewer comment #29. The fixed parameter value: $\eta = 0.1$, max. no. of iteration = 3, $w_0 = 0$, $\sigma_r^2 = 0.4$, $\sigma_x^2 = 1$, $\theta = 0.01$, $\lambda = 0.01$, $K = 33$. $K_{acq}$, $K_{ext}$, $K_{rem}$: the number of latent causes inferred during acquisition phases, extinction phases, and the remaining trials.

**Appendix 2 – table 5. Variation inflation factor of each parameter**

| Parameter | Variation inflation factor | |
|---|---|---|
| | 6-month-old group | 12-month-old group |
| $\alpha$ | 3.368 | 3.420 |
| $g$ | 2.966 | 1.472 |
| $\eta$ | 3.941 | 1.370 |
| maxIter[a] | 1.878 | 1.373 |
| $w_0$ | 2.232 | 1.644 |
| $\sigma_r^2$ | 2.292 | 1.610 |
| $\sigma_x^2$ | 2.240 | 3.343 |
| $\theta$ | 2.579 | 1.547 |
| $\lambda$ | 1.431 | 1.923 |
| $K$ | 1.612 | 2.789 |

*Note*. This table is used for the reply to reviewer comment #31. [a] max. no. of iterations.

**Appendix 2 – table 6. The joint effect of $K$, $\alpha$, and $\sigma_x^2$ on CRs (test 1 and 3), DI (test 3 vs. test 1), and the number of latent causes**

| $K$ | $\alpha$ | $\sigma_x^2$ | Test 1 | Test 3 | DI (test 3 vs test 1) | $K_{acq}$ | $K_{ext}$ | $K_{rem}$ |
|---|---|---|---|---|---|---|---|---|
| 4 | 1 | 0.01 | 1.00 | 0.34 | 0.26 | 2 | 0 | 1 |
| | 1 | 3 | 1.00 | 0.81 | 0.45 | 2 | 2 | 0 |
| | 3 | 0.01 | 1.00 | 0.35 | 0.26 | 2 | 0 | 1 |
| | 3 | 3 | 0.42 | 0.74 | 0.64 | 3 | 1 | 0 |
| 36 | 1 | 0.01 | 1.00 | 0.34 | 0.26 | 2 | 0 | 1 |
| | 1 | 3 | 1.00 | 0.34 | 0.26 | 2 | 3 | 2 |
| | 3 | 0.01 | 1.00 | 0.35 | 0.26 | 2 | 0 | 1 |
| | 3 | 3 | 0.42 | 0.19 | 0.31 | 3 | 31 | 2 |

*Note*. This table is used for the reply to reviewer comment #31. The fixed parameter value: $g$ = 1, $\eta$ = 0.1, max. no. of iteration = 3, $w_0$ = 0, $\sigma_r^2$ = 0.4, $\theta$ = 0.01, $\lambda$ = 0.01. Test 1, Test 3: the simulated CR. $K_{acq}$, $K_{ext}$, $K_{rem}$: the number of latent causes inferred during acquisition phases, extinction phases, and the remaining trials.

**Appendix 2 – table 7. Effect of $w_0$ on CR in test trials and number of latent causes**

| $w_0$ | Test 1 | Test 2 | Test 3 | $K_{acq}$ | $K_{ext}$ | $K_{rem}$ |
|---|---|---|---|---|---|---|
| -0.01 | 1 | 0.22 | 0.13 | 2 | 2 | 1 |
| -0.005 | 1 | 0.27 | 0.24 | 2 | 2 | 1 |
| 0 | 1 | 0.32 | 0.37 | 2 | 2 | 1 |
| 0.005 | 1 | 0.38 | 0.52 | 2 | 2 | 1 |
| 0.01 | 1 | 0.44 | 0.66 | 2 | 2 | 1 |

*Note*. This table is used for the reply to reviewer comment #33. The fixed parameter value: $\alpha = 1.5$, $g = 1$, $\eta = 0.1$, max. no. of iteration = 3, $\sigma_r^2 = 0.4$, $\sigma_x^2 = 1$, $\theta = 0.01$, $\lambda = 0.01$, $K = 33$. Test 1, Test 2, Test 3: the simulated CR in each test. $K_{acq}$, $K_{ext}$, $K_{rem}$: the number of latent causes inferred during acquisition phases, extinction phases, and the remaining trials.
