## Supplementary tables for "Misclassification in memory modification in *App^NL-G-F^* knock-in mouse model of Alzheimer’s disease"

**Total pages: 20**

| **Tables** | **Description** | **Reference figure or section** |
|---|---|---|
| **S1** | Cohort Table | Results, Materials and Methods section |
| **S2** | Statistical Table | Figure 2C, D, E |
| **S3, S4** | Statistical Table | Figure 2F, G, H |
| **S5, S6** | Statistical Table | Figure 3 – figure supplement 1 |
| **S7** | Statistical Table | Figure 4A |
| **S8** | Statistical Table | Figure 4B |
| **S9** | Statistical Table | Figure 4C, D, E |
| **S10** | Statistical Table | Figure 5A |
| **S11** | Statistical Table | Figure 5B |
| **S12** | Statistical Table | Figure 5C, D, E |
| **S13** | Statistical Table | Figure 6 – figure supplement 1A, B |
| **S14** | Statistical Table | Figure 6B |
| **S15** | Statistical Table | Figure 6C, D |
| **S16** | Statistical Table | Figure 6E and Figure 6 – figure supplement 1C |
| **S17** | Statistical Table | Figure 6 – figure supplement 3A, B |
| **S18** | Statistical Table | Figure 6F |
| **S19** | Statistical Table | Figure 6G, H |
| **S20** | Statistical Table | Figure 6I and Figure 6 – figure supplement 3C |
| **S21** | Statistical Table | Statistical power table |
| **S22** | Statistical Table | Statistical power table |

**Table S1. Cohort information.**

| Cohort | Group | N | Age (month old) | Sex | Instance 1 | Instance 2 |
|---|---|---|---|---|---|---|
| 1 | $App^{NL\text{-}G\text{-}F}$ | 9 | 6 | male | reinstatement | - |
| 2 | Control | 11 | 12 | male | reinstatement | - |
| 3 | $App^{NL\text{-}G\text{-}F}$ | 8 | 12 | male | reinstatement | - |
| 4[a] | Control | 25 | 6 | male | reinstatement | reversal Barnes maze |
| 5[b] | $App^{NL\text{-}G\text{-}F}$ | 17 | 6 | male | reinstatement | reversal Barnes maze |
| 6[c] | Control | 18 | 12 | male | reinstatement | reversal Barnes maze |
| 7[d] | $App^{NL\text{-}G\text{-}F}$ | 18 | 12 | male | reinstatement | reversal Barnes maze |

*Note.* The genetic background of control and $App^{NL\text{-}G\text{-}F}$ knock-in mice is C57BL/6J in cohorts 1, 2, and 3.

[a]The genotypes include C57BL/6J ($n$ = 16) and C57BL/6J-Tg(Thy1-G-CaMP7,-DsRed2) ($n$ = 9).

[b]The genotypes include C57BL/6-App<tm3(NL-G-F)Tcs>(KI/KI) ($n$ = 8) and C57BL/6J-Tg(Thy1-G-CaMP7,-DsRed2);C57BL/6-App<tm3(NL-G-F)Tcs>(KI/KI) ($n$ = 9).

[c]The genotypes include C57BL/6J ($n$ = 16) and C57BL/6J-Tg(Thy1-G-CaMP7,-DsRed2) ($n$ = 2).

[d]The genotypes include C57BL/6-App<tm3(NL-G-F)Tcs>(KI/KI) ($n$ = 14) and C57BL/6J-Tg(Thy1-G-CaMP7,-DsRed2);C57BL/6-App<tm3(NL-G-F)Tcs>(KI/KI) ($n$ = 4).

The G-CaMP7 transgeni mice (C57BL/6J-background) with and without $App^{NL\text{-}G\text{-}F}$ knock-in were well-established and used for $Ca^{2+}$ imaging (Takamura et al., 2021). When we compared the CRs for the CS between B6 and G-CaMP7 transgenic background in $App^{NL\text{-}G\text{-}F}$ knock-in and the control mice at 6 months old (cohorts 4 and 5), the magnitude relationships between the $App^{NL\text{-}G\text{-}F}$ knock-in and the control were preserved, except that only the CR in the B6 control mice was significantly lowest in test 1. Because the sample size of G-CaMP7 transgenic mice in cohort 6 was lower than 3, statistical tests for 12-month-old mice (cohorts 6 and 7) were omitted. Thus, we pooled the wild-type C57BL/6J and G-CaMP7 transgenic mice within each $App^{NL\text{-}G\text{-}F}$and control group.

### Reference

Takamura, R., Mizuta, K., Sekine, Y., Islam, T., Saito, T., Sato, M., Ohkura, M., Nakai, J., Ohshima, T., Saido, T. C., & Hayashi, Y. (2021). Modality-Specific Impairment of Hippocampal CA1 Neurons of Alzheimer's Disease Model Mice. *The Journal of Neuroscience*, *41*(24), 5315. https://doi.org/10.1523/JNEUROSCI.0208-21.2021

**Table S2. Statistical results of independent samples Student's *t*-test and one-way ANCOVA in the freezing rate during tone presentation as conditioned response (CR) in three test phases.**

| Measure | Variable | Student's *t*-test (Control vs. $App^{NL\text{-}G\text{-}F}$) | | | One-way ANCOVA | | | |
|---|---|---|---|---|---|---|---|---|
| | | $t(47)$ | *p*-value | Cohen's *d* | Effect | $F(1,95)$ | *p*-value | Partial $\eta^2$ |
| CR in test 1 phase | 6 m.o. | 2.012 | 0.05 | 0.575 | G | 9.679 | 0.002 | 0.092 |
| | 12 m.o. | 2.356 | 0.023 | 0.673 | A | 0.117 | 0.733 | 0.001 |
| CR in test 2 phase | 6 m.o. | 0.284 | 0.778 | 0.081 | G | 0.133 | 0.716 | 0.001 |
| | 12 m.o. | 0.228 | 0.82 | 0.065 | A | 4.698 | 0.033 | 0.047 |
| CR in test 3 phase | 6 m.o. | 0.601 | 0.551 | 0.172 | G | 0.561 | 0.456 | 0.006 |
| | 12 m.o. | -1.864 | 0.069 | -0.533 | A | 0.761 | 0.385 | 0.008 |

*Note.* m.o. = months old; G = Genotype (control, $App^{NL\text{-}G\text{-}F}$); A = Age (6 m.o., 12 m.o.). $n = 24$ in each control group; $n = 25$ in each $App^{NL\text{-}G\text{-}F}$ group.

**Table S3. Statistical results of one-sample and independent samples Student's *t*-test in the discrimination index (DI) for reinstatement.**

| Measure | Variable | Control | | | $App^{NL\text{-}G\text{-}F}$ | | | Student's *t*-test | | |
|---|---|---|---|---|---|---|---|---|---|---|
| | | $t(23)^a$ | *p*-value [a] | Cohen's *d* | $t(24)^a$ | *p*-value [a] | Cohen's *d* | $t(47)$ | *p*-value | Cohen's *d* |
| DI | 6 m.o. | -6.366 | <0.001 | -1.3 | -9.224 | <0.001 | -1.845 | -1.201 | 0.236 | -0.343 |
| (test 2 vs. test 1) | 12 m.o. | -4.903 | <0.001 | 0.055 | -9.812 | <0.001 | -1.233 | -2.022 | 0.049 | -0.578 |
| DI | 6 m.o. | 7.643 | <0.001 | 1.56 | 6.149 | <0.001 | 1.23 | 0.294 | 0.77 | 0.084 |
| (test 3 vs. test 2) | 12 m.o. | 5.489 | <0.001 | -1.001 | 4.32 | <0.001 | -1.962 | -1.352 | 0.183 | -0.386 |
| DI | 6 m.o. | -1.034 | 0.312 | -0.211 | -3.245 | 0.003 | -0.649 | -1.514 | 0.137 | -0.433 |
| (test 3 vs. test 1) | 12 m.o. | 0.271 | 0.789 | 1.12 | -6.165 | <0.001 | 0.864 | -3.955 | <0.001 | -1.130 |

*Note.* [a] One-sample Student's *t*-test with alternative hypothesis that the mean differs from 0.5. m.o. = months old. $n = 24$ in each control group; $n = 25$ in each $App^{NL\text{-}G\text{-}F}$ group.

**Table S4. Statistical results of one-way ANCOVA with age as a covariate in the discrimination index (DI) for the reinstatement.**

| Measure | Effect | $F(1,95)$ | *p*-value | Partial $\eta^2$ |
|---|---|---|---|---|
| DI (test 2 vs. test 1) | Genotype | 5.013 | 0.027 | 0.05 |
| | Age | 3.247 | 0.075 | 0.033 |
| DI (test 3 vs. test 2) | Genotype | 0.417 | 0.52 | 0.004 |
| | Age | 7.514 | 0.007 | 0.073 |
| DI (test 3 vs. test 1) | Genotype | 15.393 | <0.001 | 0.139 |
| | Age | 0.548 | 0.461 | 0.006 |

*Note.* Genotype effect (control, $App^{NL\text{-}G\text{-}F}$); Age effect (6 m.o., 12 m.o.). $n = 24$ in each control group; $n = 25$ in each $App^{NL\text{-}G\text{-}F}$ group.

**Table S5. Statistical results of one-sample and independent samples Student's *t*-test in the discrimination index (DI) calculated from simulated CR.**

| Measure | Variable | Control | | | $App^{NL\text{-}G\text{-}F}$ | | | Student's *t*-test | | |
|---|---|---|---|---|---|---|---|---|---|---|
| | | *t*(14 or 19)[a] | *p*-value[a] | Cohen's *d* | *t*(11 or 17)[a] | *p*-value[a] | Cohen's *d* | *t*(25 or 36) | *p*-value | Cohen's *d* |
| DI | 6 m.o. | -5.106 | < 0.001 | -1.318 | -6.713 | < 0.001 | -1.938 | -1.702 | 0.101 | -0.659 |
| (test 2 vs. test 1) | 12 m.o. | -6.4 | < 0.001 | -1.431 | -9.594 | < 0.001 | -2.261 | -2.132 | 0.04 | -0.693 |
| DI | 6 m.o. | 3.923 | 0.002 | 1.013 | 3.859 | 0.003 | 1.114 | 0.901 | 0.376 | 0.349 |
| (test 3 vs. test 2) | 12 m.o. | 3.562 | 0.002 | 0.797 | 3.173 | 0.006 | 0.748 | -0.572 | 0.571 | -0.186 |
| DI | 6 m.o. | -1.278 | 0.222 | -0.33 | -1.964 | 0.075 | -0.567 | -0.759 | 0.455 | -0.294 |
| (test 3 vs. test 1) | 12 m.o. | -1.813 | 0.086 | -0.405 | -6.271 | < 0.001 | -1.478 | -2.364 | 0.024 | -0.768 |

*Note.* [a] One-sample Student's *t*-test with alternative hypothesis that the mean differs from 0.5. m.o. = months old. $n = 15$ in 6-month-old control group; $n = 12$ in 6-month-old $App^{NL\text{-}G\text{-}F}$ group; $n = 20$ in 12-month-old control group; $n = 18$ in 12-month-old $App^{NL\text{-}G\text{-}F}$ group.

**Table S6. Statistical results of one-way ANCOVA with age as a covariate in the discrimination index (DI) calculated from simulated CR.**

| Measure | Effect | *F*(1,62) | *p*-value | Partial $\eta^2$ |
|---|---|---|---|---|
| DI (test 2 vs. test 1) | Genotype | 7.389 | 0.008 | 0.106 |
| | Age | 0.038 | 0.845 | $6.178\times10^{-4}$ |
| DI (test 3 vs. test 2) | Genotype | 0.123 | 0.727 | 0.002 |
| | Age | 4.235 | 0.044 | 0.064 |
| DI (test 3 vs. test 1) | Genotype | 5.246 | 0.025 | 0.078 |
| | Age | 3.214 | 0.078 | 0.049 |

*Note.* Genotype effect (control, $App^{NL\text{-}G\text{-}F}$); Age effect (6 m.o., 12 m.o.). $n = 15$ in 6-month-old control group; $n = 12$ in 6-month-old $App^{NL\text{-}G\text{-}F}$ group; $n = 20$ in 12-month-old control group; $n = 18$ in 12-month-old $App^{NL\text{-}G\text{-}F}$ group.

**Table S7. Statistical results of Mann-Whitney *U* test in the estimated parameters of the 12-month-old group.**

| Measure | 12 m.o. Control | | 12 m.o. *App*$^{NL\text{-}G\text{-}F}$ | | Mann-Whitney *U* | *p*-value | Effect size[a] |
|---|---|---|---|---|---|---|---|
| | Median | IQR | Median | IQR | | | |
| $\alpha$ | 1.95 | 1.72 | 1.10 | 0.28 | 100 | 0.020 | -0.444 |
| g | 1.65 | 0.90 | 1.16 | 0.49 | 151 | 0.404 | -0.161 |
| $\eta$ | 0.09 | 0.23 | 0.12 | 0.23 | 199 | 0.587 | 0.106 |
| Max. no. of iterations | 2.00 | 2.57 | 2.00 | 2.20 | 176 | 0.918 | -0.022 |
| $w_0$ | 0.006 | 0.005 | 0.002 | 0.007 | 121 | 0.087 | -0.328 |
| $\sigma_r^2$ | 1.96 | 1.78 | 1.13 | 1.50 | 145 | 0.311 | -0.194 |
| $\sigma_x^2$ | 1.57 | 1.66 | 0.35 | 0.35 | 88.5 | 0.008 | -0.508 |
| $\theta$ | 0.007 | 0.004 | 0.007 | 0.005 | 191 | 0.759 | 0.061 |
| $\lambda$ | 0.019 | 0.003 | 0.02 | 0.001 | 195 | 0.67 | 0.083 |
| K | 21.65 | 28.11 | 34.63 | 15.37 | 250.5 | 0.041 | 0.392 |

*Note.* [a]The effect size is given by the rank biserial r. $n = 20$ in control group; $n = 18$ in *App*$^{NL\text{-}G\text{-}F}$ group.

**Table S8. Spearman's rank correlation coefficient between parameters and DI in the 12-month-old group.**

| Measure | DI (test 3 vs. test 1) | *p*-value |
|---|---|---|
| $\alpha$ | 0.67 | <0.001 |
| g | 0.27 | 0.101 |
| $\eta$ | 0.13 | 0.456 |
| Max. no. of iterations | 0.03 | 0.876 |
| $w_0$ | 0.52 | <0.001 |
| $\sigma_r^2$ | 0.30 | 0.07 |
| $\sigma_x^2$ | 0.74 | <0.001 |
| $\theta$ | -0.02 | 0.897 |
| $\lambda$ | -0.03 | 0.854 |
| K | -0.73 | <0.001 |

*Note.* $n = 20$ in control group; $n = 18$ in *App*$^{NL\text{-}G\text{-}F}$ group.

**Table S9. Statistical results of Mann-Whitney *U* test in the internal state components of the 12-month-old group in test 3 phase.**

| Measure | 12 m.o. Control | | 12 m.o. $App^{NL-G-F}$ | | Mann-Whitney *U* | *p*-value | Effect size[a] |
|---|---|---|---|---|---|---|---|
| | Median | IQR | Median | IQR | | | |
| $K_{total}$ | 5 | 4 | 5 | 1.75 | 154.5 | 0.452 | -0.142 |
| Latent causes acquired in acquisition trials | | | | | | | |
| K | 3 | 1 | 2 | 0 | 101 | 0.005 | -0.439 |
| P | 0.021 | 0.02 | 0.03 | 0.022 | 191 | 0.759 | 0.061 |
| $W_{CS}$ | 0.296 | 0.52 | 0.327 | 0.334 | 202 | 0.534 | 0.122 |
| $W_{context}$ | 0.091 | 0.132 | 0.069 | 0.073 | 161 | 0.593 | -0.106 |
| Latent causes acquired in extinction trials | | | | | | | |
| K | 2 | 4 | 2 | 1 | 158 | 0.513 | -0.122 |
| P | 0.761 | 0.964 | 0.578 | 0.662 | 149 | 0.372 | -0.172 |
| $W_{CS}$ | 0.007 | 0.019 | $-3.8\times10^{-4}$ | 0.004 | 58 | 0.004 | -0.574 |
| $W_{context}$ | 0.142 | 0.164 | 0.043 | 0.091 | 84 | 0.063 | -0.382 |
| Latent causes acquired after extinction trials | | | | | | | |
| K | 0 | 1 | 1 | 0.75 | 254 | 0.02 | 0.411 |
| P | 0 | 0.431 | 0.342 | 0.444 | 243 | 0.058 | 0.35 |
| $W_{CS}$ | 0.004 | 0.003 | $-4.2\times10^{-19}$ | 0.006 | 34 | 0.145 | -0.393 |
| $W_{context}$ | 0.006 | 0.002 | 0.005 | 0.01 | 51 | 0.764 | -0.089 |

*Note.* [a]The effect size is given by the rank biserial r. $K_{total}$: total number of inferred latent causes; K: number of latent causes; P: the sum of the posterior of latent causes; $W_{CS}$: the sum of associative weight for CS; $W_{context}$: the sum of associative weight for context. $n = 20$ in control group; $n = 18$ in $App^{NL-G-F}$ group.

**Table S10. Statistical results of Mann-Whitney *U* test in the estimated parameters of the 6-month-old group.**

| Measure | 6 m.o. Control | | 6 m.o. $App^{NL\text{-}G\text{-}F}$ | | Mann-Whitney *U* | *p*-value | Effect size[a] |
|---|---|---|---|---|---|---|---|
| | Median | IQR | Median | IQR | | | |
| α | 2.50 | 1.45 | 1.13 | 1.42 | 38 | 0.012 | -0.578 |
| g | 1.86 | 1.05 | 1.10 | 1.12 | 78.5 | 0.591 | -0.128 |
| η | 0.13 | 0.65 | 0.17 | 0.72 | 99.5 | 0.66 | 0.106 |
| Max. no. of iterations | 3.75 | 2.02 | 2.00 | 2.55 | 71 | 0.366 | -0.211 |
| $w_0$ | 0.005 | 0.01 | 0.006 | 0.003 | 92.5 | 0.922 | 0.028 |
| $\sigma_r^2$ | 1.51 | 1.37 | 2.00 | 1.78 | 103.5 | 0.525 | 0.15 |
| $\sigma_x^2$ | 1.63 | 1.93 | 0.74 | 1.81 | 74 | 0.449 | -0.178 |
| θ | 0.003 | 0.004 | 0.005 | 0.008 | 145 | 0.008 | 0.611 |
| λ | 0.003 | 0.002 | 0.002 | 0.004 | 81 | 0.678 | -0.1 |
| K | 27.25 | 22.40 | 20.08 | 29.62 | 66 | 0.251 | -0.267 |

*Note.* [a]The effect size is given by the rank biserial r. *n* = 15 in control group; *n* = 12 in $App^{NL\text{-}G\text{-}F}$ group.

**Table S11. Spearman's rank correlation coefficient between parameters and DI in the 6-month-old group.**

| Measure | DI (test 3 vs. test 1) | *p*-value |
|---|---|---|
| α | 0.44 | 0.022 |
| g | 0.45 | 0.018 |
| η | 0.06 | 0.759 |
| Max. no. of iterations | 0.03 | 0.884 |
| $w_0$ | 0.13 | 0.534 |
| $\sigma_r^2$ | 0.13 | 0.533 |
| $\sigma_x^2$ | 0.43 | 0.026 |
| θ | -0.23 | 0.255 |
| λ | 0.08 | 0.701 |
| K | -0.51 | 0.006 |

*Note.* *n* = 15 in control group; *n* = 12 in $App^{NL\text{-}G\text{-}F}$ group.

**Table S12. Statistical results of Mann-Whitney *U* test in the internal state components of the 6-month-old group in test 3 phase.**

| Measure | 6 m.o. Control | | 6 m.o. $App^{NL\text{-}G\text{-}F}$ | | Mann-Whitney *U* | *p*-value | Effect size[a] |
|---|---|---|---|---|---|---|---|
| | Median | IQR | Median | IQR | | | |
| $K_{total}$ | 7 | 12.5 | 5 | 5.25 | 64.5 | 0.218 | -0.283 |
| Latent causes acquired in acquisition trials | | | | | | | |
| K | 3 | 1 | 2 | 1 | 72 | 0.321 | -0.2 |
| P | 0.015 | 0.036 | 0.033 | 0.04 | 117 | 0.196 | 0.3 |
| $W_{CS}$ | 0.127 | 0.475 | 0.156 | 0.358 | 93 | 0.905 | 0.033 |
| $W_{context}$ | 0.108 | 0.151 | 0.068 | 0.153 | 95 | 0.829 | 0.056 |
| Latent causes acquired in extinction trials | | | | | | | |
| K | 3 | 12.5 | 2.5 | 4.25 | 70.5 | 0.346 | -0.217 |
| P | 0.683 | 0.979 | 0.816 | 0.489 | 92 | 0.942 | 0.022 |
| $W_{CS}$ | 0.012 | 0.071 | 0.005 | 0.081 | 58 | 0.693 | -0.108 |
| $W_{context}$ | 0.126 | 0.095 | 0.175 | 0.201 | 75 | 0.563 | 0.154 |
| Latent causes acquired after extinction trials | | | | | | | |
| K | 0 | 1.5 | 0 | 1 | 73.5 | 0.375 | -0.183 |
| P | 0 | 0.622 | 0 | 0.268 | 75 | 0.427 | -0.167 |
| $W_{CS}$ | 0.006 | 0.008 | 0.006 | 0.003 | 15 | 0.927 | 0.071 |
| $W_{context}$ | 0.018 | 0.014 | 0.007 | 0.008 | 12 | 0.788 | -0.143 |

*Note.* [a]The effect size is given by the rank biserial r. $K_{total}$: total number of inferred latent causes; K: number of latent causes; P: the sum of the posterior of latent causes; $W_{CS}$: the sum of associative weight for CS; $W_{context}$: the sum of associative weight for context. $n = 15$ in control group; $n = 12$ in $App^{NL\text{-}G\text{-}F}$ group.

**Table S13. Statistical results of mixed-design two-way ANOVA in conventional analysis of training phases in the 12-month-old group.**

| Variable | 12 m.o. Control | | 12 m.o. $App^{NL-G-F}$ | | Effect | $F$ | df | $p$-value | Partial $\eta^2$ |
|---|---|---|---|---|---|---|---|---|---|
| | M | CI | M | CI | | | | | |
| *No. of errors* | | | | | | | | | |
| Initial training phase | | | | | | | | | |
| Day 1 | 5.82 | 1.93 | 6.19 | 3.02 | G | 0.474 | 1, 170 | 0.496 | 0.014 |
| Day 2 | 4.09 | 2.52 | 6.50 | 1.98 | D | 2.704 | 5, 170 | 0.022 | 0.074 |
| Day 3 | 4.83 | 3.02 | 5.00 | 1.80 | G × D | 1.765 | 5, 170 | 0.123 | 0.493 |
| Day 4 | 4.63 | 2.35 | 3.93 | 1.78 | | | | | |
| Day 5 | 4.43 | 3.21 | 4.43 | 2.26 | | | | | |
| Day 6 | 4.46 | 2.61 | 4.57 | 2.33 | | | | | |
| First reversal phase | | | | | | | | | |
| Day 9 | 8.54 | 3.50 | 7.85 | 4.15 | G | 0.032 | 1, 68 | 0.859 | 0.001 |
| Day 10 | 5.74 | 2.32 | 5.63 | 2.32 | D | 12.222 | 2, 68 | <0.001 | 0.264 |
| Day 11 | 4.78 | 2.22 | 5.19 | 2.08 | G × D | 0.320 | 2, 68 | 0.727 | 0.009 |
| Second reversal phase | | | | | | | | | |
| Day 12 | 4.13 | 1.24 | 4.83 | 2.78 | G | 0.136 | 1, 68 | 0.715 | 0.004 |
| Day 13 | 4.41 | 2.24 | 3.54 | 2.04 | D | 4.879 | 2, 68 | 0.010 | 0.125 |
| Day 14 | 3.20 | 2.00 | 2.76 | 2.09 | G × D | 1.390 | 2, 68 | 0.256 | 0.039 |
| *Latency* | | | | | | | | | |
| Initial training phase | | | | | | | | | |
| Day 1 | 87.59 | 31.86 | 74.50 | 32.74 | G | 6.397 | 1, 170 | 0.016 | 0.158 |
| Day 2 | 46.50 | 28.89 | 49.06 | 9.07 | D | 23.765 | 5, 170 | <0.001 | 0.411 |
| Day 3 | 40.00 | 22.36 | 30.61 | 12.63 | G × D | 2.447 | 5, 170 | 0.036 | 0.067 |
| Day 4 | 42.75 | 28.37 | 21.93 | 7.14 | | | | | |
| Day 5 | 49.28 | 40.75 | 20.02 | 8.10 | | | | | |
| Day 6 | 48.14 | 35.27 | 20.39 | 8.65 | | | | | |
| First reversal phase | | | | | | | | | |
| Day 9 | 61.56 | 25.38 | 30.30 | 18.28 | G | 10.530 | | 0.003 | 0.236 |
| Day 10 | 45.88 | 31.55 | 21.99 | 8.32 | D | 9.319 | | <0.001 | 0.215 |
| Day 11 | 41.27 | 35.29 | 18.38 | 8.46 | G × D | 0.695 | | 0.503 | 0.020 |
| Second reversal phase | | | | | | | | | |
| Day 12 | 37.03 | 30.22 | 16.54 | 7.78 | G | 10.267 | | 0.003 | 0.232 |
| Day 13 | 36.90 | 28.86 | 12.58 | 5.45 | D | 1.238 | | 0.296 | 0.035 |
| Day 14 | 34.54 | 23.77 | 11.17 | 5.63 | G × D | 0.318 | | 0.728 | 0.009 |
| *Travel distance* | | | | | | | | | |
| Initial training phase | | | | | | | | | |
| Day 1 | 416.26 | 180.53 | 403.91 | 199.55 | G | 0.278 | 1, 170 | 0.601 | 0.008 |

| | | | | | | | | | |
|---|---|---|---|---|---|---|---|---|---|
| Day 2 | 224.46 | 107.25 | 302.42 | 70.92 | D | 13.491 | 5, 170 | <0.001 | 0.284 |
| Day 3 | 239.14 | 125.57 | 221.20 | 68.53 | G × D | 1.376 | 5, 170 | 0.236 | 0.039 |
| Day 4 | 233.74 | 115.65 | 178.55 | 49.86 | | | | | |
| Day 5 | 245.30 | 194.43 | 187.16 | 72.86 | | | | | |
| Day 6 | 230.30 | 126.41 | 205.03 | 85.83 | | | | | |
| First reversal phase | | | | | | | | | |
| Day 9 | 386.83 | 149.39 | 298.62 | 148.49 | G | 1.990 | | 0.167 | 0.055 |
| Day 10 | 255.48 | 126.11 | 224.04 | 82.82 | D | 13.143 | | <0.001 | 0.279 |
| Day 11 | 242.13 | 137.53 | 206.01 | 65.90 | G × D | 0.785 | | 0.460 | 0.023 |
| Second reversal phase | | | | | | | | | |
| Day 12 | 204.57 | 81.23 | 195.45 | 72.40 | G | 1.759 | | 0.194 | 0.049 |
| Day 13 | 225.95 | 121.67 | 164.68 | 60.73 | D | 2.516 | | 0.088 | 0.069 |
| Day 14 | 183.85 | 108.40 | 147.87 | 63.82 | G × D | 1.249 | | 0.293 | 0.035 |

*Note. n* =18 in each group; G = Genotype (Control, $App^{NL\text{-}G\text{-}F}$); D = Day (1-6, 9-11, or 12-14).

**Table S14. Statistical results of Wilcoxon rank sum test in strategy analysis in Barnes maze task in the 12-month-old group.**

| Variable | z-statistics | *p*-value | Effect size r | Variable | z-statistics | *p*-value | Effect size r |
|---|---|---|---|---|---|---|---|
| *Day 1* | | | | *Day 9* | | | |
| Confirmatory | 1.01 | 0.314 | 0.17 | Confirmatory | -3.47 | 0.001 | -0.58 |
| Perimeter | 0.70 | 0.485 | 0.12 | Perimeter | 2.93 | 0.003 | 0.49 |
| Random | -1.39 | 0.163 | -0.23 | Random | -1.66 | 0.097 | -0.28 |
| Serial | -0.44 | 0.661 | -0.07 | Serial | 0.95 | 0.341 | 0.16 |
| Spatial | 0.66 | 0.512 | 0.11 | Spatial | 2.05 | 0.04 | 0.34 |
| *Day 2* | | | | *Day 10* | | | |
| Confirmatory | 1.22 | 0.221 | 0.20 | Confirmatory | -3.71 | < 0.001 | -0.62 |
| Perimeter | 1.25 | 0.211 | 0.21 | Perimeter | 2.61 | 0.009 | 0.43 |
| Random | 0.20 | 0.838 | 0.03 | Random | -0.82 | 0.411 | -0.14 |
| Serial | -3.01 | 0.003 | -0.50 | Serial | 1.06 | 0.287 | 0.18 |
| Spatial | 0.41 | 0.684 | 0.07 | Spatial | 1.74 | 0.082 | 0.29 |
| *Day 3* | | | | *Day 11* | | | |
| Confirmatory | 0.59 | 0.556 | 0.10 | Confirmatory | -2.48 | 0.013 | -0.41 |
| Perimeter | 1.80 | 0.071 | 0.30 | Perimeter | 3.10 | 0.002 | 0.52 |
| Random | -0.94 | 0.349 | -0.16 | Random | -0.24 | 0.81 | -0.04 |
| Serial | -0.67 | 0.501 | -0.11 | Serial | -2.46 | 0.014 | -0.41 |
| Spatial | 0.26 | 0.793 | 0.04 | Spatial | 0.28 | 0.778 | 0.05 |
| *Day 4* | | | | *Day 12* | | | |
| Confirmatory | 0.04 | 0.965 | 0.01 | Confirmatory | -3.77 | < 0.001 | -0.63 |
| Perimeter | 2.17 | 0.03 | 0.36 | Perimeter | 3.72 | < 0.001 | 0.62 |
| Random | -1.03 | 0.305 | -0.17 | Random | -0.50 | 0.617 | -0.08 |
| Serial | -1.55 | 0.121 | -0.26 | Serial | 0.00 | 1 | 0.00 |
| Spatial | 0.51 | 0.609 | 0.09 | Spatial | -0.19 | 0.847 | -0.03 |
| *Day 5* | | | | *Day 13* | | | |
| Confirmatory | -0.31 | 0.754 | -0.05 | Confirmatory | -2.99 | 0.003 | -0.50 |
| Perimeter | 0.95 | 0.342 | 0.16 | Perimeter | 2.09 | 0.036 | 0.35 |
| Random | 0.14 | 0.889 | 0.02 | Random | -0.47 | 0.636 | -0.08 |
| Serial | -1.37 | 0.172 | -0.23 | Serial | 2.45 | 0.014 | 0.41 |
| Spatial | 0.18 | 0.858 | 0.03 | Spatial | -0.31 | 0.757 | -0.05 |
| *Day 6* | | | | *Day 14* | | | |
| Confirmatory | -0.83 | 0.406 | -0.14 | Confirmatory | -1.87 | 0.061 | -0.31 |
| Perimeter | 0.79 | 0.427 | 0.13 | Perimeter | 2.94 | 0.003 | 0.49 |
| Random | 0.86 | 0.392 | 0.14 | Random | -0.42 | 0.676 | -0.07 |
| Serial | -0.99 | 0.321 | -0.17 | Serial | -0.83 | 0.407 | -0.14 |
| Spatial | -0.16 | 0.871 | -0.03 | Spatial | -0.25 | 0.799 | -0.04 |

*Note*. *n* =18 in each group.

**Table S15. Statistical results of mixed-design two-way ANOVA in time spent around hole in probe test of Barnes maze task in the 12-month-old group.**

| Variable | 12 m.o. Control | | 12 m.o. *App*$^{NL\text{-}G\text{-}F}$ | | Effect | *F* | df | *p*-value | Partial $\eta^2$ |
|---|---|---|---|---|---|---|---|---|---|
| | M | CI | M | CI | | | | | |
| *Probe test 1* | | | | | | | | | |
| Hole 1* | 23.35 | 6.70 | 11.00 | 2.28 | G | 3.630 | 1, 374 | 0.065 | 0.096 |
| Hole 2* | 27.84 | 11.55 | 11.83 | 2.40 | H | 9.795 | 11, 374 | <0.001 | 0.224 |
| Hole 3* | 7.07 | 2.72 | 13.98 | 9.62 | G × H | 4.223 | 11, 374 | <0.001 | 0.110 |
| Hole 4 | 4.75 | 1.69 | 8.16 | 1.25 | | | | | |
| Hole 5 | 4.16 | 1.74 | 7.34 | 2.21 | | | | | |
| Hole 6 | 3.17 | 1.62 | 5.57 | 1.39 | | | | | |
| Hole 7 | 3.46 | 2.14 | 5.86 | 1.87 | | | | | |
| Hole 8 | 3.94 | 1.86 | 5.53 | 1.45 | | | | | |
| Hole 9 | 3.11 | 1.34 | 5.09 | 1.61 | | | | | |
| Hole 10 | 4.19 | 1.76 | 5.95 | 1.84 | | | | | |
| Hole 11 | 3.43 | 1.96 | 6.58 | 1.66 | | | | | |
| Hole 12 | 13.51 | 12.43 | 7.72 | 1.75 | | | | | |
| *Probe test 2* | | | | | | | | | |
| Hole 1* | 18.35 | 5.71 | 9.74 | 1.76 | G | 0.511 | 1, 374 | 0.480 | 0.015 |
| Hole 2* | 15.77 | 6.78 | 8.16 | 1.86 | H | 4.805 | 11, 374 | <0.001 | 0.124 |
| Hole 3 | 6.46 | 2.31 | 6.99 | 1.47 | G × H | 3.242 | 11, 374 | <0.001 | 0.087 |
| Hole 4 | 4.13 | 1.85 | 7.51 | 1.74 | | | | | |
| Hole 5 | 2.51 | 1.25 | 7.56 | 1.86 | | | | | |
| Hole 6 | 3.84 | 1.26 | 7.30 | 1.54 | | | | | |
| Hole 7* | 16.34 | 13.56 | 7.80 | 1.63 | | | | | |
| Hole 8 | 6.36 | 2.11 | 5.81 | 1.05 | | | | | |
| Hole 9 | 4.05 | 1.51 | 7.48 | 1.08 | | | | | |
| Hole 10 | 5.42 | 2.22 | 7.17 | 1.73 | | | | | |
| Hole 11 | 5.72 | 1.73 | 8.09 | 1.43 | | | | | |
| Hole 12 | 7.52 | 3.60 | 9.35 | 2.55 | | | | | |

*Note. n* =18 in each group; G = Genotype (Control, *App*$^{NL\text{-}G\text{-}F}$); H = Hole (1-12). *$p < 0.05$ in Tukey's HSD test at a specific hole between control and *App*$^{NL\text{-}G\text{-}F}$ mice if a significant interaction between genotype and hole was detected.

**Table S16. Statistical results of Wilcoxon signed-rank test and Mann-Whitney *U* test in the discrimination index (DI) for Barnes maze task in the 12-month-old group.**

| Measure | 12 m.o. Control | | | | 12 m.o. $App^{NL\text{-}G\text{-}F}$ | | | | Mann-Whitney *U* | *p*-value |
|---|---|---|---|---|---|---|---|---|---|---|
| | Median | IQR | V[a] | *p*-value[a] | Median | IQR | V[a] | *p*-value[a] | | |
| DI for hole 1 | 0.50 | 0.05 | 16 | 0.477 | 0.49 | 0.22 | 70 | 0.776 | 169.5 | 0.823 |
| DI for hole 7 | 0.87 | 0.31 | 158 | 0.002 | 0.61 | 0.35 | 125 | 0.09 | 92 | 0.027 |

[a] Wilcoxon signed-rank test with alternative hypothesis that the mean differs from 0.5.

*Note.* $n$ =18 in each group. Hole 1 is the first target hole, and hole 7 is the second target hole in the reversal Barnes maze paradigm.

**Table S17. Statistical results of mixed-design two-way ANOVA in conventional analysis of training phases in Barnes maze task in the 6-month-old group.**

| Variable | 6 m.o. Control | | 6 m.o. $App^{NL\text{-}G\text{-}F}$ | | Effect | $F$ | df | $p$-value | Partial $\eta^2$ |
|---|---|---|---|---|---|---|---|---|---|
| | M | CI | M | CI | | | | | |
| *No. of errors* | | | | | | | | | |
| Initial training phase | | | | | | | | | |
| Day 1 | 6.78 | 2.00 | 7.31 | 3.20 | G | 3.434 | 1, 195 | 0.071 | 0.081 |
| Day 2 | 3.78 | 1.42 | 5.35 | 1.98 | D | 9.402 | 5, 195 | <0.001 | 0.194 |
| Day 3 | 5.13 | 2.81 | 4.53 | 1.98 | G × D | 1.160 | 5, 195 | 0.330 | 0.029 |
| Day 4 | 4.03 | 2.35 | 5.22 | 2.67 | | | | | |
| Day 5 | 2.94 | 1.51 | 4.57 | 1.84 | | | | | |
| Day 6 | 3.57 | 2.24 | 4.04 | 1.98 | | | | | |
| First reversal phase | | | | | | | | | |
| Day 9 | 6.96 | 2.38 | 6.84 | 2.26 | G | 0.023 | 1, 78 | 0.879 | 0.001 |
| Day 10 | 5.99 | 2.65 | 5.00 | 1.45 | D | 5.546 | 2, 78 | 0.006 | 0.125 |
| Day 11 | 4.49 | 2.18 | 5.35 | 2.14 | G × D | 1.147 | 2, 78 | 0.323 | 0.029 |
| Second reversal phase | | | | | | | | | |
| Day 12 | 4.61 | 1.99 | 4.73 | 2.75 | G | 0.095 | 1, 78 | 0.759 | 0.002 |
| Day 13 | 4.08 | 1.40 | 4.24 | 1.94 | D | 8.684 | 2, 78 | <0.001 | 0.182 |
| Day 14 | 3.29 | 1.43 | 2.55 | 1.14 | G × D | 0.688 | 2, 78 | 0.506 | 0.017 |
| *Latency* | | | | | | | | | |
| Initial training phase | | | | | | | | | |
| Day 1 | 88.59 | 25.02 | 93.33 | 28.64 | G | 0.041 | 1, 195 | 0.841 | 0.001 |
| Day 2 | 42.26 | 15.87 | 52.45 | 25.43 | D | 47.142 | 5, 195 | <0.001 | 0.547 |
| Day 3 | 47.54 | 40.88 | 32.01 | 15.78 | G × D | 1.474 | 5, 195 | 0.200 | 0.036 |
| Day 4 | 27.99 | 11.35 | 28.88 | 16.56 | | | | | |
| Day 5 | 21.57 | 10.18 | 23.31 | 10.22 | | | | | |
| Day 6 | 26.23 | 16.83 | 18.71 | 7.94 | | | | | |
| First reversal phase | | | | | | | | | |
| Day 9 | 42.57 | 28.66 | 33.54 | 16.47 | G | 1.836 | 1, 78 | 0.183 | 0.045 |
| Day 10 | 34.62 | 26.68 | 20.37 | 6.36 | D | 7.083 | 2, 78 | 0.001 | 0.154 |
| Day 11 | 27.41 | 19.65 | 21.25 | 8.40 | G × D | 0.577 | 2, 78 | 0.564 | 0.015 |
| Second reversal phase | | | | | | | | | |
| Day 12 | 28.18 | 23.39 | 17.56 | 8.77 | G | 1.634 | 1, 78 | 0.209 | 0.040 |
| Day 13 | 37.85 | 54.65 | 14.26 | 4.50 | D | 1.286 | 2, 78 | 0.282 | 0.032 |
| Day 14 | 22.14 | 21.18 | 12.22 | 4.61 | G × D | 0.943 | 2, 78 | 0.394 | 0.024 |
| *Travel distance* | | | | | | | | | |
| Initial training phase | | | | | | | | | |
| Day 1 | 442.45 | 112.41 | 471.44 | 136.54 | G | 0.781 | 1, 195 | 0.382 | 0.020 |

| | | | | | | | | | |
|---|---|---|---|---|---|---|---|---|---|
| Day 2 | 222.12 | 82.99 | 297.57 | 107.47 | D | 34.456 | 5, 195 | <0.001 | 0.469 |
| Day 3 | 262.03 | 115.49 | 237.77 | 121.40 | G × D | 1.192 | 5, 195 | 0.315 | 0.030 |
| Day 4 | 204.14 | 93.31 | 230.38 | 121.83 | | | | | |
| Day 5 | 158.31 | 62.00 | 197.16 | 84.05 | | | | | |
| Day 6 | 190.54 | 117.40 | 166.12 | 70.33 | | | | | |
| First reversal phase | | | | | | | | | |
| Day 9 | 289.45 | 119.65 | 267.26 | 94.01 | G | 1.051 | 1, 78 | 0.312 | 0.026 |
| Day 10 | 250.32 | 126.33 | 194.28 | 50.07 | D | 4.882 | 2, 78 | 0.010 | 0.111 |
| Day 11 | 200.42 | 78.83 | 202.05 | 56.08 | G × D | 0.645 | 2, 78 | 0.527 | 0.016 |
| Second reversal phase | | | | | | | | | |
| Day 12 | 210.36 | 99.35 | 189.77 | 84.83 | G | 1.460 | 1, 78 | 0.234 | 0.036 |
| Day 13 | 188.83 | 59.11 | 155.42 | 57.49 | D | 7.322 | 2, 78 | 0.001 | 0.158 |
| Day 14 | 153.91 | 62.07 | 129.80 | 47.29 | G × D | 0.095 | 2, 78 | 0.910 | 0.002 |

*Note.* $n$ = 24 in control group; $n$ = 17 in $App^{NL\text{-}G\text{-}F}$ group; G = Genotype (Control, $App^{NL\text{-}G\text{-}F}$); D = Day (1-6, 9-11, or 12-14).

**Table S18. Statistical results of Wilcoxon rank sum test in strategy analysis in Barnes maze task in the 6-month-old group.**

| Variable | z-statistics | *p*-value | Effect size r | Variable | z-statistics | *p*-value | Effect size r |
|---|---|---|---|---|---|---|---|
| *Day 1* | | | | *Day 9* | | | |
| Confirmatory | 0.95 | 0.341 | 0.15 | Confirmatory | 0.29 | 0.769 | 0.05 |
| Perimeter | -0.45 | 0.651 | -0.07 | Perimeter | 0.44 | 0.657 | 0.07 |
| Random | 0.03 | 0.978 | 0.00 | Random | -0.34 | 0.733 | -0.05 |
| Serial | -1.23 | 0.218 | -0.19 | Serial | 0.76 | 0.448 | 0.12 |
| Spatial | 0.21 | 0.837 | 0.03 | Spatial | -0.02 | 0.984 | -0.00 |
| *Day 2* | | | | *Day 10* | | | |
| Confirmatory | 0.43 | 0.666 | 0.07 | Confirmatory | -0.69 | 0.491 | -0.11 |
| Perimeter | 0.49 | 0.627 | 0.08 | Perimeter | 1.55 | 0.122 | 0.24 |
| Random | 1.15 | 0.250 | 0.18 | Random | -0.72 | 0.47 | -0.11 |
| Serial | -1.00 | 0.316 | -0.16 | Serial | 1.11 | 0.266 | 0.17 |
| Spatial | -0.92 | 0.355 | -0.14 | Spatial | -1.29 | 0.197 | -0.20 |
| *Day 3* | | | | *Day 11* | | | |
| Confirmatory | -0.45 | 0.651 | -0.07 | Confirmatory | -1.95 | 0.051 | -0.30 |
| Perimeter | 0.84 | 0.404 | 0.13 | Perimeter | 2.27 | 0.023 | 0.35 |
| Random | -0.54 | 0.588 | -0.08 | Random | -0.58 | 0.560 | -0.09 |
| Serial | -1.06 | 0.288 | -0.17 | Serial | -0.32 | 0.752 | -0.05 |
| Spatial | 1.40 | 0.161 | 0.22 | Spatial | 0.14 | 0.89 | 0.02 |
| *Day 4* | | | | *Day 12* | | | |
| Confirmatory | -1.96 | 0.050 | -0.31 | Confirmatory | -2.33 | 0.020 | -0.36 |
| Perimeter | 2.20 | 0.028 | 0.34 | Perimeter | 1.62 | 0.106 | 0.25 |
| Random | -0.51 | 0.611 | -0.08 | Random | -0.31 | 0.76 | -0.05 |
| Serial | -1.12 | 0.263 | -0.17 | Serial | 1.14 | 0.255 | 0.18 |
| Spatial | -0.82 | 0.411 | -0.13 | Spatial | -1.39 | 0.164 | -0.22 |
| *Day 5* | | | | *Day 13* | | | |
| Confirmatory | -0.41 | 0.681 | -0.06 | Confirmatory | -2.98 | 0.003 | -0.47 |
| Perimeter | 2.43 | 0.015 | 0.38 | Perimeter | 1.62 | 0.104 | 0.25 |
| Random | -0.50 | 0.616 | -0.08 | Random | -0.39 | 0.693 | -0.06 |
| Serial | -0.75 | 0.453 | -0.12 | Serial | 1.05 | 0.294 | 0.16 |
| Spatial | -0.37 | 0.711 | -0.06 | Spatial | -0.92 | 0.356 | -0.14 |
| *Day 6* | | | | *Day 14* | | | |
| Confirmatory | 0.00 | 1 | 0.00 | Confirmatory | -1.06 | 0.290 | -0.17 |
| Perimeter | 0.20 | 0.841 | 0.03 | Perimeter | 2.20 | 0.028 | 0.34 |
| Random | 0.43 | 0.667 | 0.07 | Random | -0.83 | 0.408 | -0.13 |
| Serial | -0.59 | 0.558 | -0.09 | Serial | -0.11 | 0.915 | -0.02 |
| Spatial | -0.03 | 0.978 | -0.00 | Spatial | -0.39 | 0.694 | -0.06 |

*Note.* $n$ = 24 in control group; $n$ = 17 in $App^{NL\text{-}G\text{-}F}$ group.

**Table S19. Statistical results of mixed-design two-way ANOVA in time spent around hole in probe test of Barnes maze task in the 6-month-old group.**

| Variable | 6 m.o. Control | | 6 m.o. *App*$^{NL\text{-}G\text{-}F}$ | | Effect | *F* | df | *p*-value | Partial $\eta^2$ |
|---|---|---|---|---|---|---|---|---|---|
| | M | CI | M | CI | | | | | |
| *Probe test 1* | | | | | | | | | |
| Hole 1 | 17.90 | 4.31 | 13.12 | 2.92 | G | 8.072 | 1, 429 | 0.007 | 0.171 |
| Hole 2 | 15.28 | 4.55 | 10.52 | 2.93 | H | 12.670 | 11, 429 | <0.001 | 0.245 |
| Hole 3 | 9.91 | 2.42 | 7.86 | 1.73 | G × H | 1.553 | 11, 429 | 0.110 | 0.038 |
| Hole 4 | 7.66 | 2.27 | 8.39 | 1.61 | | | | | |
| Hole 5 | 5.19 | 1.84 | 6.58 | 1.48 | | | | | |
| Hole 6 | 4.05 | 1.60 | 5.32 | 1.56 | | | | | |
| Hole 7 | 5.32 | 1.07 | 6.92 | 2.17 | | | | | |
| Hole 8 | 5.41 | 1.29 | 4.88 | 1.78 | | | | | |
| Hole 9 | 6.74 | 1.73 | 4.07 | 1.18 | | | | | |
| Hole 10 | 5.96 | 1.48 | 7.14 | 3.99 | | | | | |
| Hole 11 | 6.56 | 1.67 | 6.83 | 2.35 | | | | | |
| Hole 12 | 10.10 | 2.48 | 8.65 | 2.80 | | | | | |
| *Probe test 2* | | | | | | | | | |
| Hole 1* | 20.77 | 4.31 | 11.41 | 2.91 | G | 0.028 | 1, 429 | 0.867 | 0.001 |
| Hole 2 | 12.90 | 3.03 | 10.83 | 2.94 | H | 14.610 | 11, 429 | <0.001 | 0.273 |
| Hole 3 | 7.98 | 1.48 | 7.64 | 1.97 | G × H | 4.495 | 11, 429 | <0.001 | 0.103 |
| Hole 4 | 5.23 | 1.18 | 7.04 | 1.43 | | | | | |
| Hole 5 | 4.16 | 1.24 | 6.49 | 1.54 | | | | | |
| Hole 6 | 4.78 | 1.07 | 7.30 | 1.75 | | | | | |
| Hole 7 | 8.03 | 1.87 | 6.79 | 1.86 | | | | | |
| Hole 8 | 6.50 | 2.19 | 8.43 | 2.00 | | | | | |
| Hole 9 | 5.97 | 1.28 | 7.36 | 1.31 | | | | | |
| Hole 10 | 5.69 | 1.43 | 8.13 | 1.82 | | | | | |
| Hole 11 | 5.76 | 1.11 | 7.56 | 1.87 | | | | | |
| Hole 12 | 8.84 | 2.70 | 8.32 | 1.80 | | | | | |

*Note.* $n = 24$ in control group; $n = 17$ in *App*$^{NL\text{-}G\text{-}F}$ group; G = Genotype (Control, *App*$^{NL\text{-}G\text{-}F}$); H = Hole (1-12). *$p < 0.05$ in Tukey's HSD test at a specific hole between control and *App*$^{NL\text{-}G\text{-}F}$ mice if a significant interaction between genotype and hole was detected.

**Table S20. Statistical results of Wilcoxon signed-rank test and Mann-Whitney *U* test in the discrimination index (DI) for Barnes maze task in the 6-month-old group.**

| Measure | 6 m.o. Control | | | | 6 m.o. $App^{NL\text{-}G\text{-}F}$ | | | | Mann-Whitney *U* | *p*-value |
|---|---|---|---|---|---|---|---|---|---|---|
| | Median | IQR | V[a] | *p*-value[a] | Median | IQR | V[a] | *p*-value[a] | | |
| DI for hole 1 | 0.51 | 0.09 | 120 | 0.042 | 0.46 | 0.14 | 45 | 0.410 | 134 | 0.064 |
| DI for hole 7 | 0.68 | 0.35 | 226 | 0.029 | 0.43 | 0.35 | 88.5 | 0.586 | 169 | 0.365 |

[a] Wilcoxon signed-rank test with alternative hypothesis that the mean differs from 0.5.

*Note.* $n = 24$ in control group; $n = 17$ in $App^{NL\text{-}G\text{-}F}$ group. Hole 1 is the first target hole, and hole 7 is the second target hole in the reversal Barnes maze paradigm.

**Table S21. The statistical power reference table (results in Figure 1 to 5).**

| Test type | N1 | N2 | Effect size metric | power | | | | |
|---|---|---|---|---|---|---|---|---|
| | | | | 0.1 | 0.3 | 0.5 | 0.7 | 0.9 |
| *CR, DI* | | | | | | | | |
| independent samples Student's t test | 24 | 25 | Cohen's *d* | 0.190 | 0.419 | 0.572 | 0.725 | 0.946 |
| independent samples Student's t test | 15 | 12 | Cohen's *d* | 0.263 | 0.578 | 0.789 | 1.001 | 1.307 |
| independent samples Student's t test | 20 | 18 | Cohen's d | 0.218 | 0.479 | 0.654 | 0.829 | 1.082 |
| *DI in each group* | | | | | | | | |
| one-sample Student's t-test | 24 | - | Cohen's *d* | 0.139 | 0.306 | 0.418 | 0.529 | 0.691 |
| one-sample Student's t-test | 25 | - | Cohen's *d* | 0.136 | 0.299 | 0.408 | 0.518 | 0.676 |
| one-sample Student's t-test | 15 | - | Cohen's *d* | 0.181 | 0.397 | 0.543 | 0.690 | 0.901 |
| one-sample Student's t-test | 12 | - | Cohen's *d* | 0.206 | 0.453 | 0.620 | 0.787 | 1.029 |
| one-sample Student's t-test | 20 | - | Cohen's *d* | 0.154 | 0.338 | 0.462 | 0.585 | 0.764 |
| one-sample Student's t-test | 18 | - | Cohen's *d* | 0.163 | 0.358 | 0.490 | 0.621 | 0.811 |
| *CR, DI* | | | | | | | | |
| one-way ANCOVA | 24 | 25 | Partial $\eta^2$ | 0.009 | 0.042 | 0.076 | 0.116 | 0.183 |
| one-way ANCOVA | 15 | 12 | Partial $\eta^2$ | 0.017 | 0.076 | 0.134 | 0.199 | 0.297 |
| one-way ANCOVA | 20 | 18 | Partial $\eta^2$ | 0.012 | 0.054 | 0.097 | 0.147 | 0.226 |
| *Estimated parameters, internal states* | | | | | | | | |
| Mann-Whitney U test | 15 | 12 | r | 0.140 | 0.297 | 0.391 | 0.475 | 0.576 |
| Mann-Whitney U test | 20 | 18 | r | 0.117 | 0.250 | 0.333 | 0.409 | 0.505 |
| *Correlation between parameters and DI* | | | | | | | | |
| Spearman's rank order regression | 15 | 12 | ρ for H1 | 0.132 | 0.281 | 0.374 | 0.460 | 0.571 |
| Spearman's rank order regression | 20 | 18 | ρ for H1 | 0.109 | 0.236 | 0.316 | 0.392 | 0.493 |
| Spearman's rank order regression | 15 | - | P for H1 | 0.184 | 0.384 | 0.499 | 0.599 | 0.717 |
| Spearman's rank order regression | 12 | - | ρ for H1 | 0.212 | 0.433 | 0.557 | 0.659 | 0.774 |
| Spearman's rank order regression | 20 | - | ρ for H1 | 0.156 | 0.329 | 0.434 | 0.528 | 0.644 |
| Spearman's rank order regression | 18 | - | ρ for H1 | 0.166 | 0.348 | 0.457 | 0.553 | 0.671 |

*Note.* The level of $\alpha$ is 0.05. Rank-biserial $r = d/(d^2+4)^{1/2}$, where d is Cohen's *d* calculated in G*Power. Partial $\eta^2 = f^2/(f^2+1)$, where f is Cohen's *f* calculated in G*Power.

**Table S22. The statistical power reference table (results in Figure 6).**

| Test type | N1 | N2 | Effect size metric | power | | | | |
|---|---|---|---|---|---|---|---|---|
| | | | | 0.1 | 0.3 | 0.5 | 0.7 | 0.9 |
| *Day 1 to Day 6* | | | | | | | | |
| mixed design two-way ANOVA | 24 | 17 | Partial $\eta^2$ | 0.006 | 0.030 | 0.054 | 0.085 | 0.136 |
| mixed design two-way ANOVA | 18 | 18 | Partial $\eta^2$ | 0.007 | 0.034 | 0.062 | 0.096 | 0.153 |
| *Day 9 to Day 11 or Day 12 to Day 14* | | | | | | | | |
| mixed design two-way ANOVA | 24 | 17 | Partial $\eta^2$ | 0.007 | 0.034 | 0.062 | 0.095 | 0.152 |
| mixed design two-way ANOVA | 18 | 18 | Partial $\eta^2$ | 0.008 | 0.039 | 0.070 | 0.108 | 0.171 |
| *12 holes in Probe test* | | | | | | | | |
| mixed design two-way ANOVA | 24 | 17 | Partial $\eta^2$ | 0.006 | 0.028 | 0.051 | 0.079 | 0.127 |
| mixed design two-way ANOVA | 18 | 18 | Partial $\eta^2$ | 0.007 | 0.032 | 0.058 | 0.090 | 0.144 |
| *Strategy analysis, DI* | | | | | | | | |
| Wilcoxon rank-sum test | 24 | 17 | r | 0.112 | 0.241 | 0.322 | 0.396 | 0.490 |
| Wilcoxon rank-sum test | 18 | 18 | r | 0.120 | 0.257 | 0.342 | 0.419 | 0.516 |
| *DI in each group* | | | | | | | | |
| Wilcoxon signed-rank test | 24 | - | r | 0.075 | 0.163 | 0.221 | 0.276 | 0.351 |
| Wilcoxon signed-rank test | 17 | - | r | 0.091 | 0.197 | 0.265 | 0.330 | 0.415 |
| Wilcoxon signed-rank test | 18 | - | r | 0.088 | 0.191 | 0.257 | 0.320 | 0.404 |

*Note.* The level of $\alpha$ is 0.05. Partial $\eta^2 = f^2/(f^2+1)$, where f is Cohen's *f* calculated in G*Power. $r = d/(d^2+4)^{1/2}$, where d is Cohen's *d* calculated in G*Power.
